# Neural mechanisms underlying the temporal control of sequential saccade planning in the frontal eye fields

**DOI:** 10.1101/2020.12.04.411454

**Authors:** Debaleena Basu, Naveen Sendhilnathan, Aditya Murthy

## Abstract

Sequences of saccadic eye movements are instrumental in navigating our visual environment. While neural activity has been shown to ramp up to a threshold before single saccades, the neural underpinnings of multiple saccades is unknown. To understand the neural control of rapid saccade sequences, we recorded from the frontal eye field (FEF) of macaque monkeys while they performed a sequential saccade task. We show that concurrent planning of two saccade plans brings forth processing bottlenecks, specifically by decreasing the growth rate and increasing the threshold of saccade-related ramping activity. The rate disruption affected both saccade plans, and a computational model wherein activity related to the two saccade plans bilaterally and asymmetrically inhibited each other, predicted the behavioral and neural results observed experimentally. Borrowing from models in psychology, our results demonstrate a capacity-sharing mechanism of processing bottlenecks, wherein multiple saccade plans in a sequence, compete for the processing capacity by perturbation of the saccade-related ramping activity. Finally, we show that in contrast to movement related neurons, visual activity in FEF neurons is not affected by the presence of multiple saccade targets, indicating that for perceptually simple tasks, inhibition amongst movement-related neurons mainly instantiates capacity sharing. Taken together, we show how psychology-inspired models of capacity sharing can be mapped onto neural responses to understand the control of rapid saccade sequences.

## Introduction

Saccadic eye movements shift the fovea from one point to another, serially sampling our visual surroundings, and aiding consequent behavior. Proper planning and execution of saccade sequences is essential for performing everyday tasks such as reading. Despite extensive research on the neural basis of planning individual saccades, the neural mechanisms underlying the sequencing of multiple saccades remain largely unknown. Previous research has shown that sequential saccades can be processed in parallel (Basu and Murthy, 2020; Becker and Jürgens, 1979; Bhutani et al., 2012; Bhutani et al., 2013; McPeek et al., 2003; McPeek and Keller, 2002; McPeek et al., 2000; Minken et al., 1993; Phillips and Segraves, 2010; Port and Wurtz, 2003; Ray et al., 2004; Sharika et al., 2008; Shen and Paré, 2014; Tian et al., 2000; Wu et al., 2013). Sequential saccade studies have shown that as the temporal gap between the targets (TSD; target step delay) decreases, the latency of the response to the second stimulus increases markedly, as if the brain inherently cannot process two simple decisions at the same time (Pashler, 1994; Marois and Ivanoff, 2005; Ray et al., 2012; Ray et al., 2004; Ruthruff et al., 2001). The bottlenecks associated with parallel programming of multiple saccade plans form the basis of this study.

Various theoretical frameworks have been proposed to explain how closely spaced action plans interfere with each other. Single-channel bottleneck models propose that a central, decision-making stage constitutes the bottleneck, wherein the central stages of multiple plans can only proceed serially and cannot be ‘co-active’ (Pashler, 1994; Ruthruff et al., 2001; Welford, 1967; Welford, 1952). For a sequence of two saccades, the first plan is likely to reach the central stage first, and thus the saccade 2 plan must ‘wait’ till central processing of the first is over (**Fig. 1A)**. In contrast, capacity-sharing models argue that the decision-making stages of both plans can proceed in parallel, albeit with differential rates. The concept of the brain’s ‘capacity’ corresponds to the brain’s general information processing capabilities (Broadbent, 1971; Gopher and Navon, 1980; Kahneman, 1973; McLeod, 1977), independent of task type. The capacity-sharing models predict that because of its temporal precedence, the first saccade plan will get the major share of the capacity and the second saccade plan will get a smaller fraction, thus delaying the onset of the second response (**Fig. 1B**; Navon and Miller, 2002; Tombu and Jolicœur, 2003).

**Figure 1.**
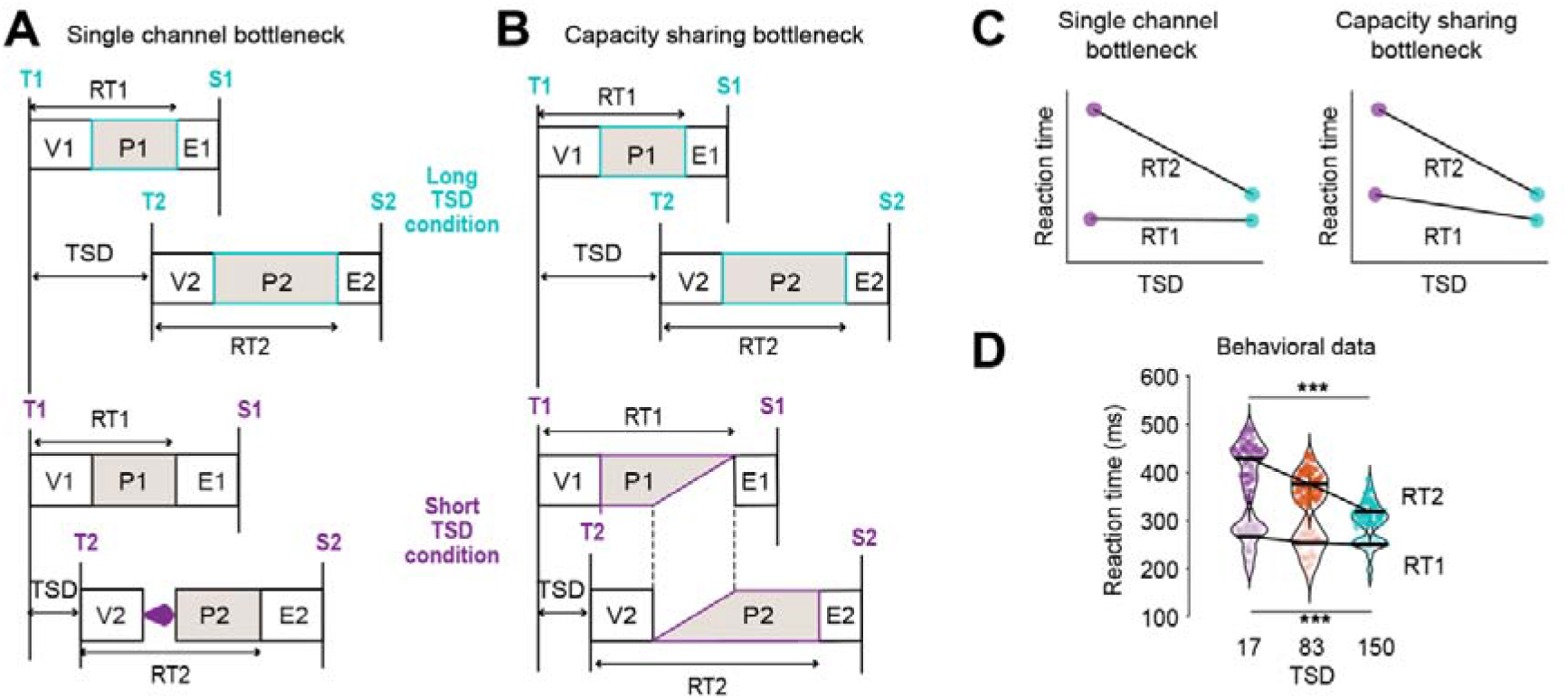
Behavioral predictions for processing bottlenecks during the planning of sequential saccades. **A.** Single-channel bottleneck framework. Each task is made up of three stages. The visual stage (V) can be carried on in parallel with stages of another task, but the central planning stage, P, can only proceed singly. In a two-saccade sequence, the stages of the first saccade plan proceed to completion unabated leading to its execution (E). For the second plan however, if the second target closely follows the first (low TSD condition), the central planning stage, P2, is postponed till P1 is complete. Such a postponement does not occur in the long TSD condition, where the two saccade plans are well-separated, thereby leading to an increase of RT2 from long to short TSD. **B.** Capacity-sharing bottleneck framework. In this framework, the P stages of multiple plans can proceed in parallel and access the brain’s limited processing capacity simultaneously. In the low TSD condition, P1 and P2 concurrently ‘share’ the capacity, resulting in slower progress of the saccade plans. This leads to lengthening of both RT1 and RT2 in the low TSD condition, the effect on RT2 being greater as the second saccade plan gets a smaller share of the central capacity. **C.** Predictions of reaction time vs TSD for single-channel bottleneck framework (left) and capacity-sharing bottleneck framework (right). RT2 increases with decrease in TSD for both frameworks, whereas RT1 increase is predicted only by the capacity-sharing model. **D.** Behavioral data for reaction time vs TSD. Data shows trials in which the first (for RT1) or second (for RT2) saccade was into the response field. Both reaction times increased significantly with decrease in TSD.

The neural mechanisms of processing bottlenecks in sequential saccade planning are not known. To investigate the neural architecture of saccade-related bottlenecks, we recorded neural activity from the frontal eye field (FEF) of macaque monkeys performing a sequential saccade task. FEF is a good candidate region to study the neural imprints of processing bottlenecks since it is a higher-order control center for goal-directed saccadic planning (Sendhilnathan et al., 2021; Sendhilnathan et al., 2017, 2020). Further, the activity of FEF movement neurons follow the dynamics of accumulator models and resemble the central capacity-limited stage observed in computational models of dual-task studies (Hanes and Schall, 1996; Ray et al., 2012; Sigman and Dehaene, 2005). Finally, FEF movement neurons can encode two saccade plans in parallel (Basu and Murthy, 2020), and thus, any limitations arising during the concurrent programming of saccades may be found in the activity of movement-related neurons in the FEF. Our results show that FEF movement neurons constitute a bottleneck locus—the processing of saccadic sequences is slowed down by reducing the speed of activity growth or by increasing movement activation threshold. Such adjustments were observed for both the first and second saccade plans, indicating that a capacity-sharing mechanism might underlie temporal delays seen during the sequencing of multiple actions.

## Results

Two monkeys, a *Macaca radiata* (J) and a *Macaca mulatta* (G) performed a sequential saccade ‘FOLLOW’ task (**Fig. S1**; see methods), where the majority (70%) of the trials were ‘step trials’ in which they had to perform a rapid sequence of saccades to two targets in the order of their presentation. The remaining 30% of the trials were ‘no-step’ trials, wherein a single visual target was presented, and the monkeys had to make a single saccade to it. The two types of trials were randomly interleaved. The temporal gap (target step delay or TSD) between the first and second target onsets in step trials was randomly chosen among 17 ms, 83 ms, and 150 ms (Basu and Murthy, 2020).

### Behavioral evidence of processing bottlenecks during sequential saccades

In the scheme of single-channel bottleneck models, the second plan shows the hallmark of processing bottlenecks: increase in latencies with decrease in TSD, whilst the saccade 1 latencies (RT1) stay unaffected (**Fig 1C** left). However, unlike the single-channel bottleneck model, where plan 1 may be assumed to get 100% of the capacity, the capacity-sharing model predict that the latencies of the first saccade (RT1) will also increase as it only gets a part of the full available capacity (**Fig 1C** right).

To ensure that the behavioral data are matched to the neural data, we analyzed trials in which saccades were made into the response field (RF; see methods). That is, for RT1, the first saccade was made into the RF, and for RT2, the second saccade was made into the RF. Both RT1 and RT2 slowed down as the TSD decreased, indicating a capacity-sharing mechanism (**Fig. 1D**; RT1: Kruskal-Wallis, *χ^2^* (2, 240) = 17.85, *p* < .001, η^2^ = 0.07; RT2: Kruskal-Wallis, *χ^2^* (2, 233) = 158.37, *p* < .001, η^2^ = 0.67). While the effect on RT1 was typically much smaller than that on RT2, the increases in saccade latencies with decreasing TSD corroborated with previously well-established evidence of processing bottlenecks in concurrent action planning. Our behavioral data, thus, supports the presence of a capacity-sharing bottleneck as opposed to the single-channel bottleneck as the first saccade plan does not stay unaffected.

### Movement-related activity during single saccades

Previous work has shown that the pattern of activity of FEF movement neurons are correlated with stochastic accumulation, which is widely used in computational models of saccadic reaction times (Boucher et al., 2007; Hanes and Schall, 1996; Ratcliff et al., 2007; Woodman et al., 2008) and are directly linked to saccade initiation times (Huerta et al., 1986; Langer and Kaneko, 1990; Segraves, 1992). Since reaction time lengthening is the main behavioral evidence of processing bottlenecks, movement neurons are well projected to carry neural correlates of the same. To confirm whether movement-related activity in FEF adheres to an accumulation-to-threshold model of reaction time, we first studied the no-step single-saccade trials. We divided these trials into fast, medium, and slow reaction time groups and we measured the parameters of accumulator models from the movement activity (**Fig. S2A**). The reaction time grouping was obtained by partitioning reaction times in each session using the mean reaction time of that session (see methods). The main parameters of accumulator models, i.e., baseline, onset, growth rate, and threshold activity were measured for the three-reaction time conditions (fast, medium, slow), for each neuron (**Fig. S2C-F;** see methods). Consistent with the earlier studies (Hanes and Schall, 1996), adjustments in the rate of growth of activity of the movement neuron population predicted reaction times in the no-step trials: across the movement neuron population, the slope of the best fitting line for growth rate variation in the reaction time groups was significantly different from zero (Z_rate_ = −4.27, p < .001; **Fig. S2E**). Further, the slopes for the growth rate were negative, indicating that fast reaction times were preceded by a steeper rate of growth of movement activity and vice versa. While the growth rate varied with reaction time, the threshold did not (Z_threshold_ = −0.98, p = .323; **Fig. S2F**), corroborating with the established reaction time models of accumulation to a fixed threshold. The slope distributions of other accumulator measures like baseline, and onset, were not statistically significant from zero (Z_baseline_ = −2.04, p = 0.05; Z_onset_ = 1.92, p = 0.054).

### Processing bottlenecks underlie the representation of sequential saccades

Using a computational model, Sigman and Dehaene (2005) had shown that evidence accumulation, representing a central decision process, constituted a bottleneck in dual-tasks, while the perceptual stage and the execution stage did not. Based on the mapping between accumulator models and movement neuron dynamics, four possible hypotheses (**Fig. S2A**) can explain how the activity of FEF movement neurons coding for the second saccade might bring about the systematic increase of the latency of the second saccade (RT2) with decrease in TSD that characterizes processing bottlenecks. The lengthening of reaction time may be due to (1) lowering of the baseline firing rate with shorter TSDs (2) delaying of the onset of the activity related to the second saccade with shorter TSDs (3) reduced growth rate of the activity with shorter TSDs (4) and an increase of the saccade threshold firing rate with larger TSDs.

**Fig. 2** schematically shows the possible modulations of the accumulation process in the planning stage (P) and the corresponding movement neuron activity. The accumulation process is represented as a noisy integrator accumulating visual evidence till it reaches the threshold. In the single-channel bottleneck model, RT1 is unaffected, and thus the dynamics of the integrator and the corresponding neural activity will be unchanged across the three TSDs (**Fig. 2A**). For RT2, the single-channel bottleneck model posits a postponement of the central stage, thus the onset of the accumulating process and of the neural activity will get delayed as the overlap between the two saccade plans increases from long to short TSD (**Fig. 2B**). According to the capacity-sharing model, the first and second saccade plans can proceed in parallel; thus, there is no ‘waiting period’ for the accumulation process of the second plan—the onset of neural activity will be similar across TSDs for both first and second saccade. However, since both motor plans share the limited processing capacity, the central stages of both plans will be lengthened. This may be brought about by a decrease in the rate of integration from long to short TSD, or an increase in the decision threshold. At the level of neural activity, the rate of ramping up of movement-related activity may slow down, or the threshold firing rate for saccade onset may increase to account for the increase in saccade latencies with decrease in TSD. Critically, the rate and/or threshold modulation will be present for both saccade plans according to the capacity-sharing model, although the effect may be lesser for the first plan as the corresponding increase in RT1 is also less (**Fig. 2C** & **Fig. 2D**). While we have presented polarized scenarios for the two bottleneck models, it is possible that at the population level, there would a combination of the factors mentioned.

**Figure 2.**
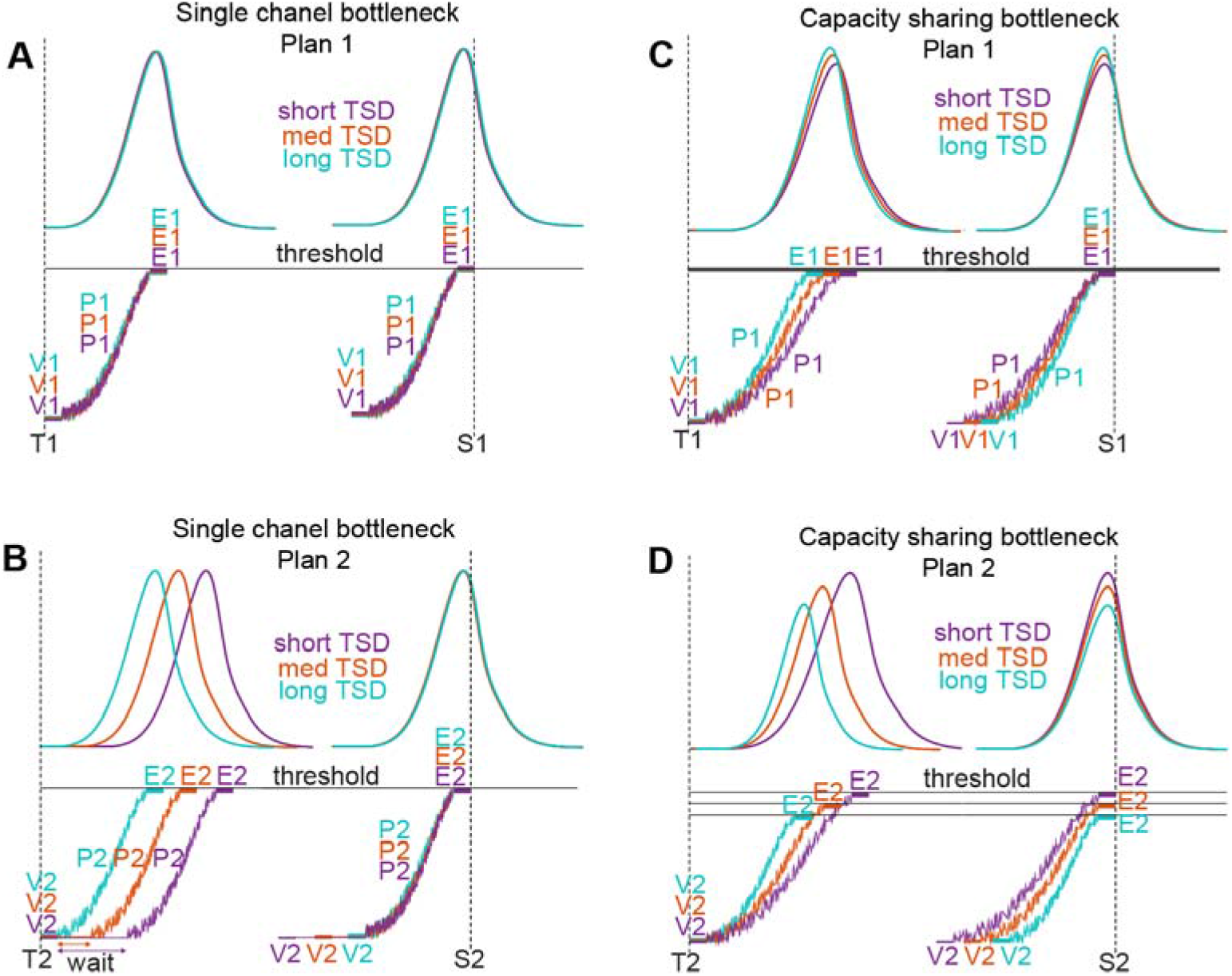
Neural activity predictions for processing bottlenecks during the planning of sequential saccades. **A.** Hypothesized neural activity for saccade plan 1 in the single-channel bottleneck framework: Bottom: After the first visual target is presented (vertical broken line; T1), there is an initial visual processing stage (V1) which is often of constant duration for all plans. The planning stage (P1) for the first saccade is represented as a noisy integrator, accumulating activity till the motor threshold (horizontal solid line) is reached and the saccade is executed (E1). Top panel: The corresponding neural activity is shown as the ramping up of FEF movement neuron activity till saccade onset (S1). The activities corresponding to three different TSDs are shown in three different colors. **B.** Hypothesized neural activity for saccade plan 2 in the single-channel bottleneck framework. The onset of the accumulation process and the ramping up of the neural activity will shift later with decrease in TSD to account for RT2 elongation (same format as A; T2: onset of second target; V2: visual processing stage for target 2, P2: planning stage for saccade 2; E2: execution stage for plan 2; S2: onset of second saccade). **C.** Hypothesized neural activity for saccade plan 1 in the capacity-sharing bottleneck framework. The onsets of the integrators and the movement neuron activity do not change with TSD on account of parallel programming of the two saccade plans. Increase in saccade latencies at shorter TSDs maybe brought about by a decrease in the growth rate from long to short TSD. Same format as **A**. **D.** Hypothesized neural activity for saccade plan 2 in the capacity-sharing bottleneck framework. Same as C, with the addition of threshold modulation and a greater degree of rate adjustment with TSD to constitute the larger increase in RT2 from long to short TSD. Same format as **A**.

To assess which of the above possibilities explain the increase in RT2, we analyzed the neural activity in trials where the second saccade was made into the RF (RFin trials; see methods) for all three TSDs (**Fig. 3A**; see **Fig S3** for single neuron example). Across the population, the rate of neural activity growth slowed down from long to short TSD, and the activity ramped up to a higher firing rate threshold. We measured each of the four accumulator parameters: baseline, onset, rate, and threshold (averaged across trials of the same TSD) for the three TSD conditions, for each neuron (**Fig. 3B;** see methods) using linear regression. The slopes from all the movement neurons were compared using a Wilcoxon signed-rank test. Across the movement neuron population, the slopes of the rate and the threshold, as a function of TSD were significantly different from zero (Z_rate_ = 4.27, p < .001; Z_threshold_ = −2.67, p < 0.01; **Fig. 3B**). Further, the slopes for the rate of activity growth were positive, indicating that the rate of activity grew faster at longer TSDs, where presumably the effect of processing bottlenecks was the least among the three TSD conditions. Threshold slopes were significantly negative, indicating that as the TSD increased, the threshold required for initiation of the second saccade was reduced at the population level. However, the slope distributions of other accumulator measures like baseline, and onset, were not statistically significant from zero (Z_baseline_ = −0.62, p = 0.53; Z_onset_ = 0.86, p = 0.17). Thus, processing bottlenecks at the level of FEF movement neurons were characterized by multifaceted adjustments in the rate and threshold of the activity related to S2.

**Figure 3.**
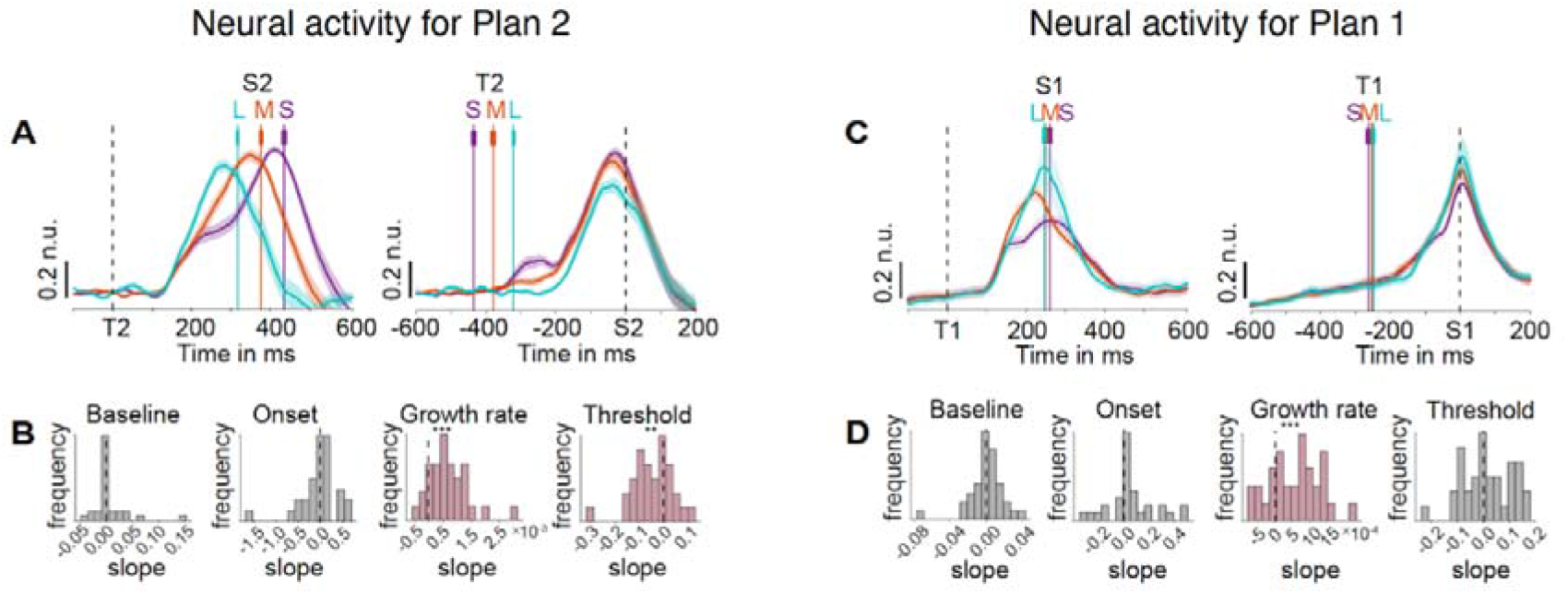
Activity for FEF movement neurons during sequential saccades. **A. Top:** Population activity of FEF movement neurons when the second saccade went into the response field, aligned on second target onset (T2). Mean saccade onset times (S2) for the short, medium, and long TSD conditions are shown as vertical, colored lines with s.e.m error bars. Right: same as left but activity aligned to the second saccade onset. Shading indicates mean ± SEM. **B.** Population histogram of slopes of each measure of accumulator dynamics (baseline, onset, growth rate and threshold) as a function of TSD for FEF movement neurons. Asterisks denote cases where the distribution of movement neuron slopes was significantly different from zero (Wilcoxon signed-rank test, *** *p* < .001, ** *p* < .01). **C.** Same as **A** but for saccade 1 **D.** Same as **B** but for saccade 1

While the classical evidence of processing bottlenecks is indexed by the increase in RT2, RT1 may also be affected according to the capacity-sharing scheme of processing bottlenecks (**Fig. 1B**). We tested whether movement-related activity encoding first saccade remained unchanged as would be expected in the single-channel bottleneck scheme, or changed systematically, across TSDs as the capacity-sharing model predicted. To address this issue, we performed the same analyses as before but for the condition in which the first saccade was made into the RF (RFout trials; **Fig 3C**; see **Fig S3** for single neuron example). At the population level, rate perturbation occurred with decrease in TSD in the first plan, mirroring the modulation observed for the second plan (Wilcoxon signed-rank test for slopes of rates, Z_rate_ = 3.62, p < .001; **Fig 3D**). However, unlike the second plan, threshold activity did not show a significantly decreasing relation with TSD (Z_threshold_ = 1.16, p = 0.25;). Slope distributions of other accumulator measures like baseline, and onset, were not statistically significant from zero (Z_baseline_ = 0.85, p = 0.39; Z_onset_ = 1.04, p = 0.29; **Fig 3D**). Thus, rate perturbation constituted a major mechanism through which the ramping up of activity of FEF movement neurons was controlled during parallel planning of sequential saccades.

### State space dynamics and inhibitory control may enable capacity sharing during sequential saccade planning

To gain deeper insights into neural mechanisms underlying capacity sharing, we studied the population dynamics underlying the trajectory of neural activity in FEF. First, we visualized this by performing a principal component analysis (PCA) separately for the population neural activity (for saccades into RF) aligned to target 1 and target 2 onsets for each of the three TSDs (**Fig 4A**). PCA is a commonly used unsupervised learning algorithm to extract the latent information from the data (see methods). This method allows us to look at the high dimensional FEF population neural activity in a much lower dimension that captures the maximum variance of the population. At least ~ 7-8 PCs were required to explain >99% of the variance for any of the six conditions (three TSDs, two plans; although there was a trend of fewer PCs explaining more variance as TSD increased). However, the top three PCs explain >90% variance. Therefore, we visualized a ‘state-space trajectory’ by plotting the top three PCs versus one another (**Fig 4B**). Each point on the trajectory indicates the neural state at each time point.

**Figure 4:**
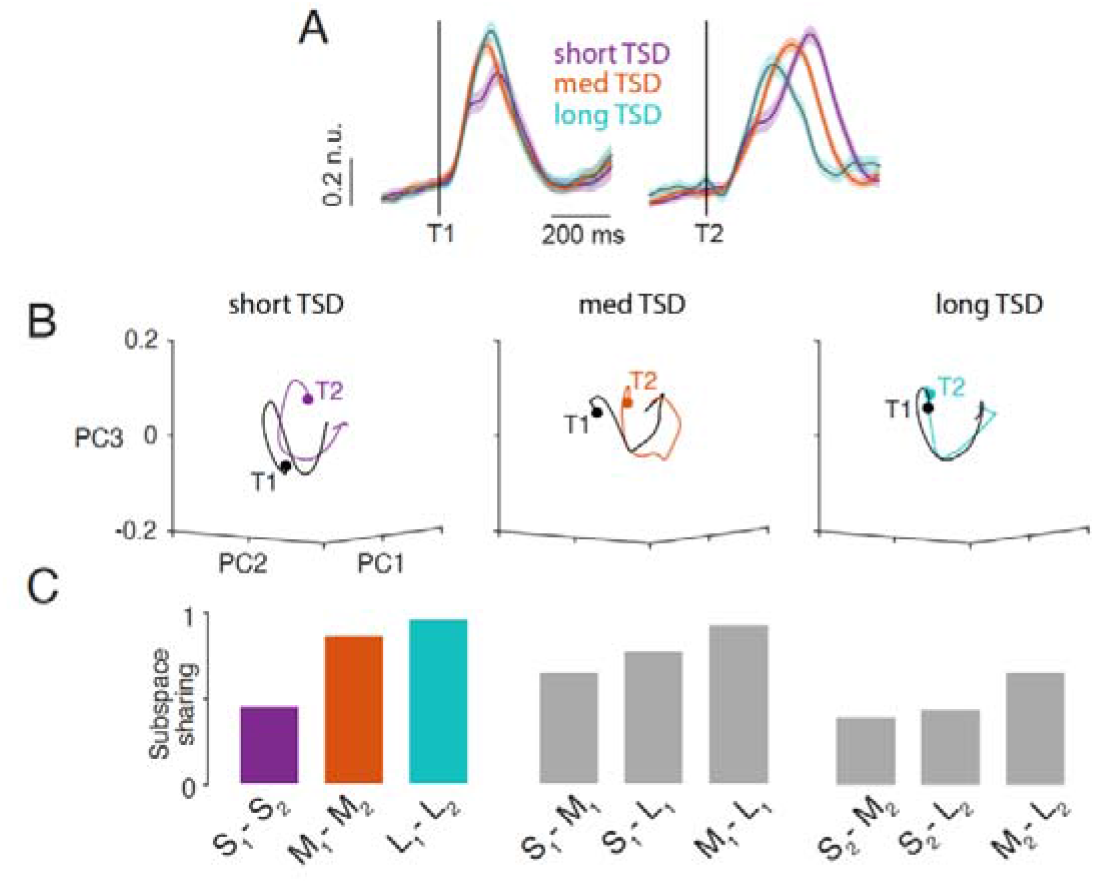
Extent of subspace sharing explains processing bottlenecks during the planning of sequential saccades. **A.** Normalized mean population neural responses aligned to target 1 (T1) and target 2 (T2) for short, medium and long TSD trials (n.u. = normalized unit); same as **Fig 3A** and **3C.** **B.** Cumulative percent variance explained by the first 10 PCs for target (T1) (black) and target 2 (T2) (color) related responses for short (left), medium (center) and long (right) TSD trials. **C.** Subspace overlap between a pair of conditions. S, M and L indicate short, medium and long TSDs, and 1 and 2 indicate the saccade plan number.

If planning for the first and the second saccades are processed in parallel but compete for the same shared space due to limited capacity (according to the capacity sharing model), we should expect the neural trajectories to span different subspaces at shorter TSDs and span the same subspace at higher TSDs. That is, in the lowest possible TSD, we should expect the two subspaces to be completely orthogonal (no overlap) and as the TSD increases and approaches the reaction time of the first saccade, the subspaces can begin to overlap. Therefore, in our case with the lowest TSD being 17 ms, we should expect a low degree of overlap and at TSD = 150 ms (~RT1), we should expect a high degree of overlap. In contrast, the single-channel bottleneck hypothesis predicts that the subspaces corresponding to the planning of the first and the second saccades would completely overlap, since the plan 2 would be completely dormant until plan 1 is completed.

We found that the neural trajectories significantly differed between the planning of the first and the second saccades for the shortest TSD but became more similar as the TSD increased (**Fig 5D**). We quantified the degree of overlap between the subspaces spanned by these neural trajectories (see methods). At the shortest TSD, the magnitude of overlap between the signals for planning of the first and the second saccades was 47% and this increased as the TSD increased from medium (84%) to long TSDs (92%; **Fig 4C**), aligned more with the predictions of the capacity sharing model. This result also held true for all saccade directions (**Fig S4**). We also confirmed that these differences were related to the TSDs and not to differences in saccade kinematics, which were similar across TSDs for the first and the second saccades (**Fig S5**).

**Figure 5:**
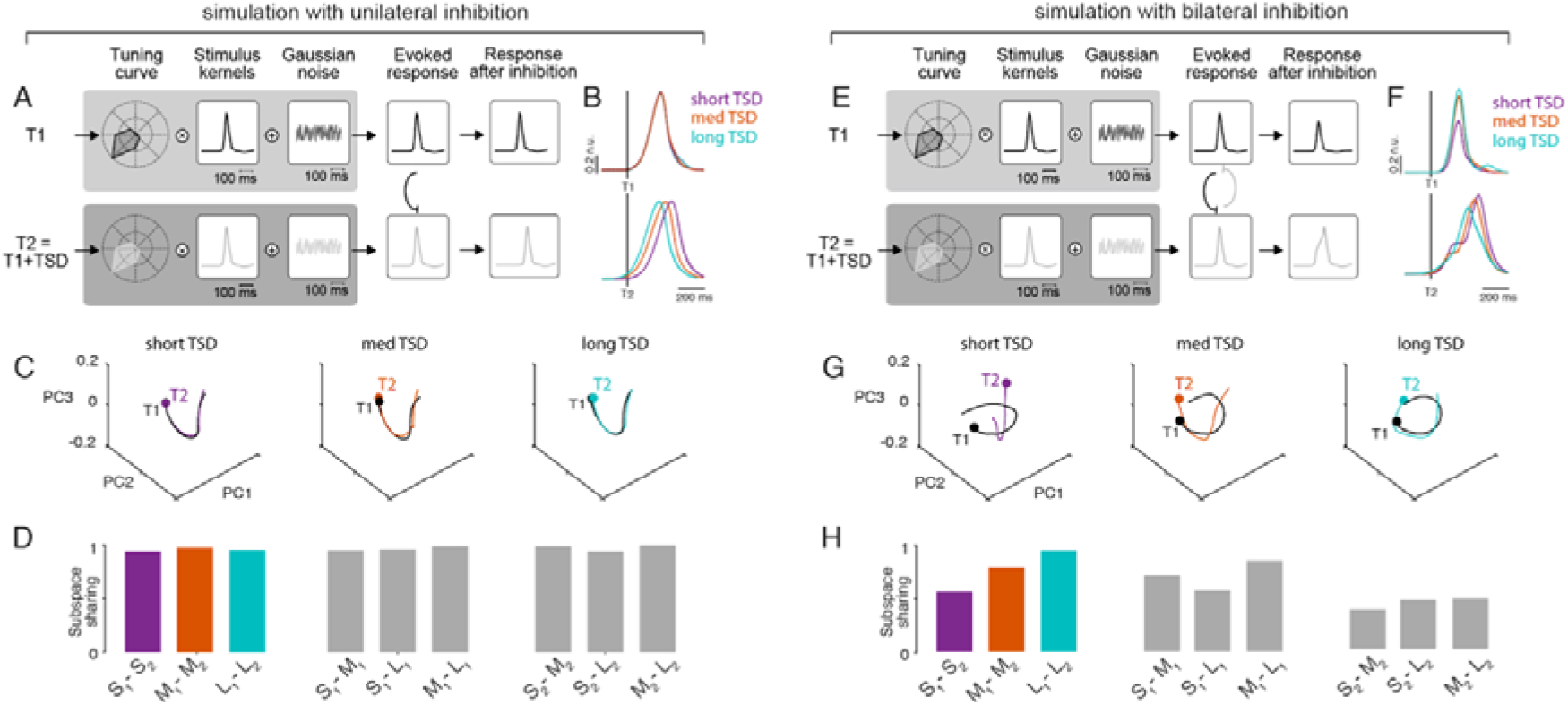
Only simulations with bilateral, asymmetric inhibition capture the empirical data’s population dynamics. **A.** Schematic of the simulation with bilateral inhibition. Top row: simulated neural activity for the first saccade plan and bottom row: simulated neural activity for the second saccade plan (see **Fig S6** and methods; n.u. =normalized unit). **B.** Normalized mean population neural responses, for data simulated with bilateral inhibition, aligned to target 1 and target 2 for short, medium and long TSD trials. **C.** First three PCs plotted against each other for target 1 (black) and target 2 (color) related responses for short (left), medium (center) and long (right) TSD trials. Filled circle markers indicate the starts of the respective trajectories. **D.** Subspace overlap between a pair of conditions. S, M and L indicate short, medium and long TSDs, and 1 and 2 indicate the plan number. **E.** Same as **A,** but for simulation with unilateral inhibition (see methods). **F.** Same as **B**, but for simulated data with unilateral inhibition. **G.** Same as **C**, but for simulated data with unilateral inhibition. **H.** Same as **D**, but for simulated data with unilateral inhibition.

Next, we investigated the mechanism behind the differences in the neural subspace overlap among different TSDs. We performed two sets of simulations (see methods; **S5A-F**) using the accumulator framework (**Fig S5H**). For each of the two sets, we simulated 40 neurons with 900 trials per neuron (with three types of TSD trials) using a firing rate model to approximately match the statistical power of our experimental dataset (see methods). We constructed an inhibition function such that the magnitude of the inhibition inversely varied with TSD (see methods; **Fig S5G**).

In the first set of simulations, we introduced a unilateral inhibition (**Fig 5A;** see methods). Here, the activities for plan 2 were temporally shifted by plan 1 following the inhibition curve as a function of TSD. The resulting simulated neural activities (**Fig 5B**) resembled the predictions of a single-channel bottleneck model (**Fig 2A-B**). Very few (~3) PCs explained >99% of the variance. The state-space neural trajectories were not significantly different between planning of the first and the second saccades for any of the TSDs (**Fig 5C)** as the subspace overlap was 98% between any pair of plans (**Fig 5D**), as expected from the single-channel bottleneck model.

In the next set of simulations, we introduced bilateral, asymmetric mutual inhibition (see methods; **Fig 5E**). That is plan 1 temporally shifted plan 2 just like before but plan 2 reduced the magnitude of peak firing of plan 1. Hence the nature of inhibition is both bilateral and asymmetric. The simulations of this model (**Fig 5F**) resembled the neural data (**Fig 3A, C**) and the predictions of a capacity sharing bottleneck model (**Fig 2C-D**). Here, ~ 7-8 PCs were required to explain >99% of the variance for the shortest TSD and fewer (~5-6) PCs were required to explain >99% of the variance for the longest TSD. The neural trajectories significantly differed for short TSD but were similar for longer TSDs (**Fig 5G**) and the degree of subspace overlap between the two plans increased with TSD, consistent with the structure present in the neural data (**Fig 5H**) resembling the experimental data (**Fig 4C**), as expected from the capacity sharing model.

### FEF visually-related neurons do not show processing bottlenecks

Previous studies have reported a separation between the visual and motor processing of FEF neurons with only motor processing affecting reaction time in perceptually simple tasks (Sato et al., 2001; Thompson et al., 1997; Woodman et al., 2008). Thus, it is plausible that the responses of visual neurons are not gated by inhibitory bottlenecks. This notion was tested by analyzing target-related activity in purely visual (**Fig 6A**) and visuomovement neurons (**Fig 6B**).

**Figure 6.**
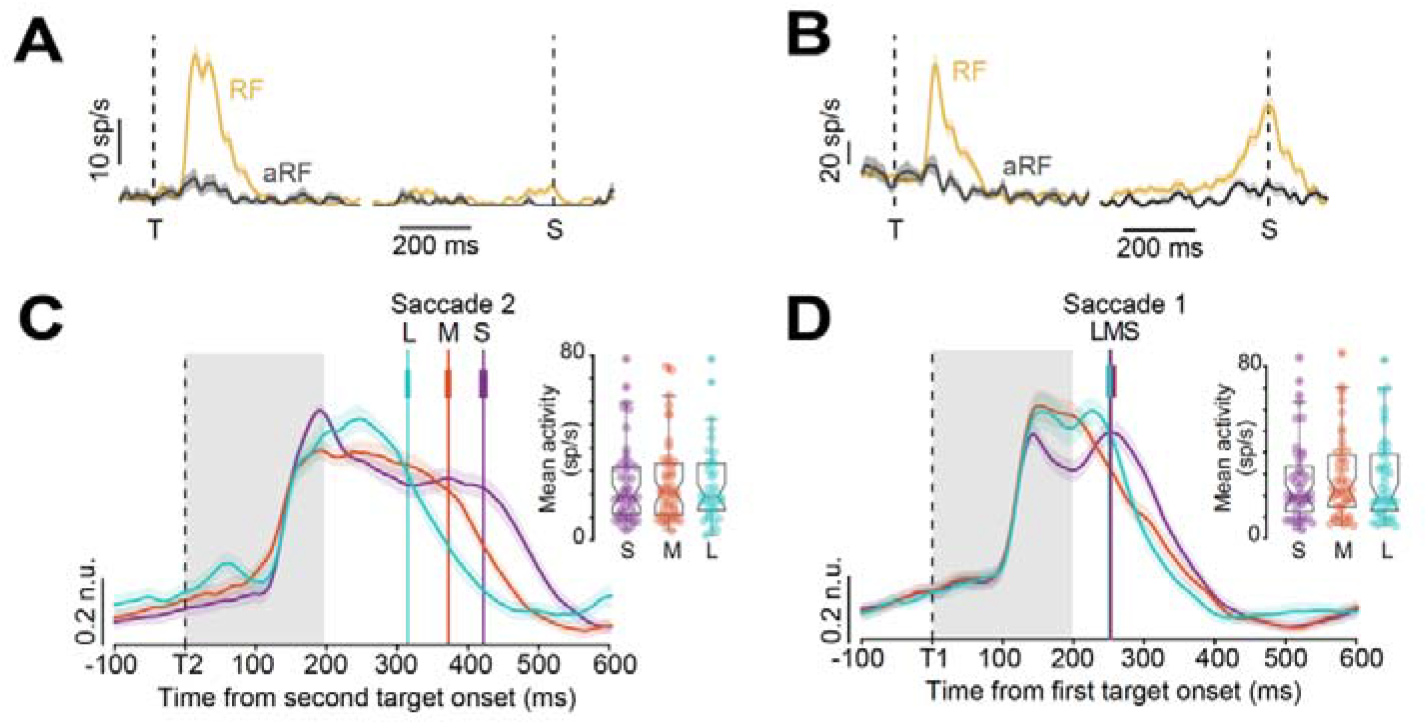
Processing bottlenecks in FEF V and VM neurons. **A.** A representative FEF visual neuron aligned to target onset (T) and saccade onset (S) in a memory-guided task for saccades into the RF (yellow) and saccades out of RF (into aRF; black). **B.** A representative FEF vismov neuron aligned to target onset (T) and saccade onset (S) in a memory-guided task for saccades into the RF (yellow) and saccades out of RF (into aRF; black). **C.** FEF visual and vismov neuron population activity encoding for different TSD conditions (short, medium, long), aligned to target 2 onset. Mean saccade onset times for the TSD conditions are shown as vertical colored lines with s.e.m error bars. Inset: Kruskal–Wallis box plots for average activity in the visual epoch (gray shaded area) for the three TSDs. Shading indicates mean ± SEM. **D.** Same as **C** but for neural activity aligned to target 1.

We analyzed the average target-related response in the 200 ms window following target onset for each neuron to identify signatures of processing bottlenecks. If target selection is capacity-limited, then presumably, neural responses encoding saccade targets appearing in close succession will be inhibited, either due to single-channel bottleneck (only second target response gets affected) or due to capacity sharing (both first and second target responses get affected). In contrast to movement-related activity, the average activity in the target-related period did not vary with TSD (Kruskalwallis: *χ^2^* (2, 129) = 0.47, *p* = .79 (first saccade); *χ^2^* (2, 124) = 0.06, *p* = .97 (S2)) for both saccade plans, suggesting that the visual processing stage is pre-bottleneck, at least of a perceptually simple task like the FOLLOW task.

## Discussion

In this study, we explored the limits of parallel processing involved in saccade sequences. Processing bottlenecks were found within FEF, the mechanisms being rate perturbation and threshold modulation in the movement neuron population. Additionally, we found evidence of processing bottlenecks for both motor plans for the first and the second saccades, suggesting that the associated bottleneck could be a consequence of capacity sharing between co-activated movement plans. The notion of such shared and limited processing was also revealed in the state space dynamics of FEF movement activity, which showed a potential role for inhibitory control that gated access of concurrent motor plans to a planning subspace. Our analysis of visual activity did not reflect any consistent modulation that could be considered a significant bottleneck. The major results are discussed and interpreted in the following sections.

### Processing bottlenecks in sequential saccade planning

Processing bottlenecks and parallel programming represent functionally antithetical processes, and yet both are essential for optimal saccadic behavior. While parallel programming allows for rapid execution of a saccade sequence, processing bottlenecks are likely to arise to check unbridled parallel programming of motor plans, as failure to control it might lead to errors like averaged saccades or incorrect order of execution of a saccade sequence (Bhutani et al., 2012; Coëffé and O’regan, 1987; Findlay, 1982; Ray et al., 2012; Viviani and Swensson, 1982; Zambarbieri et al., 1987). In the context of the current study, we tested whether a single-channel bottleneck (Pashler, 1994) or a capacity-sharing bottleneck (Kahneman, 1973) best explained our reaction time data since behavioral evidence of both the models have been found in dual-task paradigms (Arnell and Duncan, 2002; Navon and Miller, 2002; Pashler, 1994). In our data, we found evidence of increase in both RT1 and RT2 with TSD, ruling out the single-channel bottleneck model being the exclusive framework underlying bottlenecks in sequential saccades. Our neural data also suggested a capacity-sharing mechanism of bottlenecks: the onset of saccade-related activity did not vary with TSD as predicted by the single-channel bottleneck hypothesis (see **Fig. 2**), and both saccade plans showed consistent activity modulations with TSD. A reduction in the rate of accumulation and an increase in the threshold activity level were seen for the second saccade plan. In contrast, only changes in the slope of the activity corresponding to the first saccade were observed, which may account for the more subtle changes in RT1.

### Inhibitory control underlying processing bottlenecks

We tested whether mutually inhibitory accumulators encoding distinct saccade plans can mimic capacity sharing, wherein both the saccadic eye movements are executed with delays, especially for the second saccade. Modelling such a response required two important conditions: the first condition required that the inhibition be asymmetric, being greater for the first saccade plan than the second saccade plan, which manifest as greater capacity and faster information processing for the former compared to the latter. Such an asymmetry is a natural consequence of the temporal delay allowing for greater activity in the first saccade to inhibit the second saccade; the second condition required an inhibitory kernel that decreased with target step delay, such that inhibition from the first accumulator to be greater at shorter delays despite being the level of activity in the accumulator being lesser compared to what it would be at larger target step delays. Such an inhibitory kernel is necessary to match the observed behavioral data of greater second saccade reaction times as well as the neural data which showed greater interference for the second saccade motor plans at the shorter TSDs. Interestingly, using a dynamical systems approach under the assumption of stationarity of noise across trials (Elsayed et al., 2016), this model of inhibitory control could be also shown to act as a “queuing” mechanism, in which non-orthogonal neural spaces can simultaneously allow parallel processing but yet temporarily slow the processing of the second saccade. We believe that the ability of such inhibition to reconfigure the neural space may reflect the nonlinear effects of inhibition on the pattern of activity representing accumulator activity that underlie the saccades.

The simplest and most parsimonious explanation for the location of such a bottleneck is at the level of FEF via bilateral mutual inhibition (Ray et al., 2009) of competing motor plans developing in the FEF. This type of inhibitory gating can be brought about by inhibitory interneurons within the FEF (Markram et al., 2004; Somogyi, 1977). Although such a form of inhibition is intuitive and can be readily implemented within the proposed frameworks described for decision-making circuits (Bogacz et al., 2006; Ratcliff and Smith, 2004), implementing an inhibitory kernel that decreases with increasing TSD cannot be easily implemented in a straightforward manner by mutually inhibitory accumulators. Furthermore, using an identical task, our previous work has shown that the basal ganglia is causally involved in the conversion of parallel movement plans into sequential behavior (Bhutani et al., 2013). Inactivation of the basal ganglia in monkeys with muscimol or impairment of the basal ganglia in Parkinson’s disease patients resulted in a significantly greater extent of saccadic errors that develop due to unchecked parallel programming leading to a ‘collision’ of saccade plans. The results of both these studies can be reconciled by the fact that FEF and basal ganglia share a closed connection through the cortico-BG-thalamo-cortical loop wherein the thalamus, a major relay center, receives projections from BG output nuclei, and in turn projects to multiple cortical regions, including the FEF, which are again routed to the input nuclei of basal ganglia (Alexander et al., 1986; Middleton and Strick, 2000; Parent and Hazrati, 1995a, b). Thus, the origin of the bottleneck could also be in the well-established inhibitory control circuitry of the basal ganglia (Hikosaka et al., 2000) and then re-routed to the FEF through the basal ganglia-thalamo-cortical loop (Goldman-Rakic and Porrino, 1985), which then manifests it various adjustments of movement-related neuronal activity.

### Neural representations of processing bottlenecks within FEF

Our data show robust signatures of processing bottlenecks involving rate and threshold adjustments of FEF movement neurons contributing to the observed processing bottlenecks. Interestingly, similar adjustments of rate have been observed in FEF movement-related neurons when monkeys slow their reaction times to improve their accuracy (Heitz and Schall, 2012), consistent with movement-related activity reflecting a developing motor plan that can be adjusted by strategic requirements of the task. However, in contrast to speed/accuracy adjustments, we did find systematic increases in threshold for the second saccade with shorter TSDs that together with decreases in accumulation rate, contribute to the lengthening of reaction times for the second saccade. Interestingly, similar changes in growth rate for both the first and second saccade, particularly at shorter TSDs, were also observed in our model of mutually inhibiting accumulators but without any changes in threshold (**Fig 5**), raising the possibility that these changes may involve additional processes such as adjustments in the excitability of superior colliculus neurons from the basal ganglia (Lo and Wang, 2006; Wurtz and Hikosaka, 1986) that were not modelled here.

In contrast to the movement neurons, the activity of visual neurons displayed little evidence of active inhibitory control, suggesting that they are ‘pre-bottleneck’. This is not surprising since many studies have reported a separation between the visual and motor processing of FEF neurons with only motor processing affecting reaction time in perceptually simple tasks, thus it is plausible that the responses of visual neurons are not gated by inhibitory bottlenecks for our task. However, it can be speculated that in a more perceptually challenging task, manifestations of processing bottlenecks would show up in the activity of visual responses as well. Thereby, it can be concluded that movement neurons, which are thought to be functionally downstream of visual neurons (Woodman et al., 2008), are subjected to a greater degree of inhibitory control, possibly due to its direct role in saccade initiation. A similar result was observed in the countermanding (Hanes et al., 1998) and redirect tasks (Murthy et al., 2009), where movement-related neurons showed the strongest evidence of inhibitory control that reflected the monkeys’ abilities to withhold or change saccade plans. Thus, movement-related activity would fall under the ‘post-/peri-bottleneck’ category while visually-related activity would be ‘pre-bottleneck’, at least for perceptually simple tasks.

## METHODS

### KEY RESOURCES TABLE

| REAGENT OR RESOURCE | SOURCE | IDENTIFIER |
| --- | --- | --- |
| Experimental models: Organisms/Strains |  |  |
| Rhesus macaque (*Macaca mulatta*) |  | N/A |
| Bonnet macaque (*Macaca radiata*) |  | N/A |
| Software and Algorithms |  |  |
| MATLAB | Mathworks | https://www.mathworks.com/products/matlab.html |
| Blackrock | Blackrock Microsystems | https://www.blackrockmicro.com/ |
| ISCAN eye tracking system | ISCAN | http://iscaninc.com/ |
| Others |  |  |
| Tungsten microelectrode | FHC | https://www.fh-co.com/product/metal-microelectrodes/ |

### CONTACT FOR REAGENT AND RESOURCE SHARING

Further information and requests for resources and reagents should be directed to and will be fulfilled by the Lead Contact, Debaleena Basu.

### EXPERIMENTAL MODEL AND SUBJECT DETAILS

The detailed methods pertaining to this dataset has been published in a previous study (Basu & Murthy 2020; Sendhilnathan et al., 2021). A brief overview is given below.

#### Experimental Animals

Single-unit recordings were done from two adult monkeys (J, male *Macaca radiata*, and G, female *Macaca mulatta*). The animals were cared for in accordance with the animal ethics guidelines of the Committee for the Purpose of Control and Supervision of Experiments on Animals (CPCSEA), Government of India, and the Institutional Animal Ethics Committee (IAEC) of the Indian Institute of Science (IISc.).

#### Surgical Procedures

Each monkey underwent two surgeries: first, to implant a titanium headpost for the purpose of head-fixation during experiments, and second to make an MRI-guided craniotomy over the FEF and implant a recording chamber (Crist instruments, USA). Training or recording sessions were conducted only after the monkeys completed surgical recovery.

### METHOD DETAILS

#### Behavioral tasks

Monkeys were trained on two oculomotor tasks: the memory-guided (MG) saccade task and the FOLLOW saccade task. Trials in the MG task began with a red fixation point (0.6°× 0.6°) which was presented in the center of a screen. After a variable fixation period (~300 ms), a gray target stimulus (1°× 1°) was presented peripherally. Post-appearance, the target disappeared after 100 ms; however, the monkeys were required to fixate for a delay period of around 1000 ms. The fixation spot was extinguished after the delay period, following which the monkeys had to make a saccade to the remembered target location. Correct trials were reinforced with juice rewards. The delay period served to aid the classification of FEF neurons by isolating the stimulus-related (visual) and saccade-related (motor) epochs.

The FOLLOW task (**Fig S1**) is a modified version of the double-step task (Becker and Jürgens, 1979; Westheimer, 1954; Wheeless et al., 1966), where single saccade no-step trials (30%) were randomly interleaved with sequential saccade step trials (70%). Trials started with fixation, following which a green saccade target (1°× 1°) was presented in one of the six possible peripheral locations (eccentricity 12°). The fixation spot was removed at target onset. In no-step trials, the monkeys had to execute a single, correct saccade to the target. In step trials a red second target (1°× 1°) appeared after target 1, signaling the monkey to make ordered sequential saccades. Step trials comprised two targets, the first one being same as in the no-step trials. After a variable time delay (target step delay (TSD): 17 ms, 83 ms, or 150 ms), a red target was displayed. Monkeys had to make an additional second saccade from target 1 to target 2 to get rewarded in step trials.

Response field (RF) identification was done using the MG task. The RF center, and the two flanking positions were set as ‘RFin’ positions and the three diametrically opposite positions were considered ‘RFout’ positions. No-step targets and the first target of step trials could appear at any one of the six RFin and RFout locations. The second target in step trials was presented in any one of three positions diametrically opposite to the location of the first target. Based on this scheme, RFin trials refer to trials in which the second target or target 2 was presented in the RF while target 1 was outside RF. RFout trials are those in which target 1 was inside RF and target 2 was outside. Neural activity in RFin trials would mainly encode the second target or second saccade, while RFout trials would represent the first target or the saccade.

#### Recording setup and procedures

The tasks were designed and displayed using TEMPO and VIDEOSYNC software (Reflective computing, St. Louis, MO, USA). A Sony Bravia LCD monitor (42 inches, 60 Hz refresh rate; 640 × 480 resolution) was used to show the task stimuli to the monkeys. An infrared eye tracker (ISCAN, Woburn, MA USA) was used to track the pupils throughout the recording session.

Neural recordings were undertaken using tungsten microelectrodes (FHC, Bowdoin, ME, USA, impedance 2 - 4 MΩ). A Cerebus data acquisition system (Blackrock Microsystems, Salt Lake City, UT, USA), which was synchronized to the TEMPO software, was used to sample and store neuronal data at 30,000 Hz. In each recording session, the MG task was used to identify and classify FEF neurons. After RF and cell type was identified, the FOLLOW task was started.

### QUANTIFICATION AND STATISTICAL ANALYSIS

#### Data Analysis

The collected neural data was sorted offline using the in-built spike-sorting tool of Cerebus system (Blackrock Microsystems). Saccades were detected from eye position data using a 30°/s velocity threshold. Analysis of the data was done using MATLAB (MathWorks, Natick, MA, USA). This study used only trials with correct responses, with at least eight trials per condition as the inclusion criteria. The final dataset for this study comprised 84 FEF neurons. A filter mimicking an excitatory post-synaptic potential (EPSP) was used to convolve spike data (Murthy et al., 2007).

The classification of FEF neurons was done using the MG task. The delay period separated the visual epoch (90-180 ms after target onset) from the movement epoch (80 ms window preceding saccade onset). Visual neurons were identified if the activity was increased in the visual epoch compared to baseline (300-100 ms preceding target onset), movement neuron displayed higher activity in the movement epoch, and visuomovement neurons showed increased activity in both the epochs. A visuo-motor index (VMI) was used to validate the cell classification (Murthy et al., 2007).

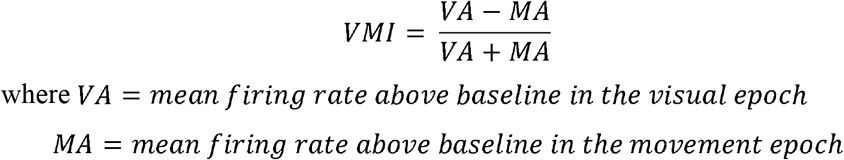

With the range being from +1 to −1, visual neurons had positive VMIs, while movement neurons had negative VMIs. Visuomovement neurons yielded intermediate VMIs. Activity in the single saccade trials of the FOLLOW task was also taken in account for proper cell classification, as changing task contexts have been shown to influence neuronal activity profiles (Jagadisan and Gandhi, 2016).

#### Accumulator parameters

Taking cue from accumulator models and previous studies (Woodman et al., 2008), four main parameters were calculated: (1) Baseline firing rate; (2) Onset of firing rate increase; (3) Threshold activity required for saccade initiation; (4) Rate of growth of activity from onset to threshold. These measures of accumulator dynamics were calculated separately for FOLLOW step trials in which the first saccade went into the RF (RFout) and those in which the second saccade was towards the RF (RFin). Since correct FOLLOW step trials always had a sequence of two saccades stepping from an RF-in position to an RF-out position or vice versa, the activity in the RFin trials was a mix of RFout and RFin activities. To specifically analyze activity that contributed only to the second saccades made into the RF, the mean RF-out activity of no-step trials was used as a reference and subtracted from the mean activity in RFin step-trials. For single neurons, the parameter calculations were made from non-normalized, differential activity for RFin trials.

For FOLLOW no-step trials, trials in which the saccade was into the response field were used for accumulator parameter calculation. Trials in each session were grouped into fast, medium, or slow reaction time trials based on the average reaction time of that session, i.e. trials with reaction time less than 30 ms below mean reaction time were considered as fast trials, those with reaction time more than 30 ms above mean reaction time were slow trials, and trials around the mean reaction time (± 10 ms) were medium reaction time trials. Accumulator parameters were then calculated for the three-reaction time groups.

Baseline activity was measured as the average of the differential activity in the RFin condition in the 100 ms before the appearance of the first FOLLOW target. Onset was defined as the time point when FEF activity first exceeded 2 SDs above baseline, provided that the differential activity ultimately reached 4 SDs and was maintained above 2 SDs for at least 50 ms for the second saccade plan and 20 ms for the first saccade plan. Threshold activation was the average firing rate in the RF-in condition in the interval from 10 to 20 ms before saccade initiation (Hanes and Schall, 1996). Rate of activity growth was measured by subtracting the threshold-activity level from the onset-activity level and dividing by the time interval between onset and threshold. This measure was robust against fluctuations in the rise-profile. To better understand non-linear rise profiles, the rate was also measured by piecewise regression fits using a sliding window of width 40 ms (for RFin trials) from onset to threshold and calculating the slopes and intercepts at each point. For population analyses, difference SDFs of each session were normalized to the peak average activity in the TSD = 17 ms group and for each saccade plan i.e. activity related to second saccade plan was normalized with respect to TSD 17 activity for second saccades going into the RF and vice versa for the first saccade plan. In the case of no-step trials, the SDFs were normalized to the peak activity of the fast trials in each session.

#### Principal Components Analysis (PCA)

For analyses based on dimensionality reduction, we performed two steps of data preprocessing before further analyses. First, we ‘soft normalized’ the neural responses (*r*) for each neuron, *i*, by dividing the neural activity for each neuron (*r_i_*) by its range (|*r_i_*| =*r_i_*/(*range*(*r_i_*))) (Churchland et al., 2012). Soft normalization preserves the structure of inter-neuronal variation while normalizing the population response so that neurons with strong responses could be reduced to approximately unity range, but neurons with weak responses could be reduced to less than unity range. Second, we mean-centered the responses of each neuron by subtracting the mean activity of a given neuron across all conditions 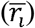 from the neural response 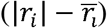.

To identify the signals that best represent the population activity of neurons, we performed principal component analysis (PCA), a common unsupervised learning algorithm, on the data. To do this, we constructed two matrices P_1_ and P_2_ of size *t* × *n* where *t* is the time and n is the number of neurons with population response for the first and the second saccade plans, respectively. We applied PCA to P_1_ and P_2_ yielding W_1_ and W_2_ respectively, which are *n* × *k* matrix each, of principal components.

We used a metric to index the degree to which the population response occupied different neural dimensions on trials with different TSDs. To compute this ‘subspace overlap’ between the first and the second saccade plans for each TSD, we first defined the variance captured as 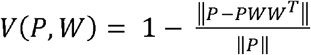, where the operator ∥X∥ means the Frobenius norm of the matrix X. Then, the subspace overlap was given by: 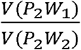.

Subspace overlap should be equal to one if the population responses occupied the same dimensions (i.e., are spanned by the same PCs) on both the saccade plans. And the subspace overlap should be equal to zero if the population responses occupied mutually orthogonal dimensions on both the saccade plans.

#### Neural Simulations

We simulated 40 motor neurons with 900 trials (with three types of TSD trials) using a firing rate model (**Fig S5A**) to approximately match the statistical power of our experimental dataset. We defined the spatial properties of each neuron (tuning curve) through a cosine function centered on one of the 8 positions which was randomly chosen and called it the neuron’s RF:

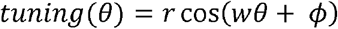

where *r* is the peak firing rate, *w* defines the width of the tuning curve, *θ* is a set of 8 target positions and *ϕ* is the displacement (**Fig S5B**). We then defined the temporal properties of the neurons (**Fig S5C**) using a skewed Gaussian distribution response kernel:

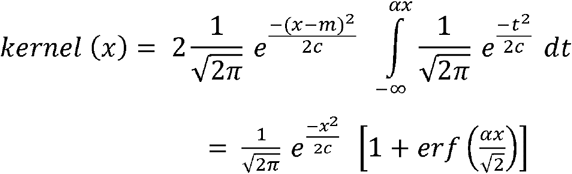

where *c* = 8000 controls the full width at half maximum of the distribution, *erf*(*x*) is the error function and *α* is the shape parameter that controls the shape of the distribution. To simulate a noisy accumulator (**Fig S5H**), we sampled 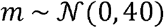 and 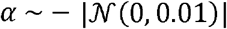. Note that similar results can be obtained using a response kernel resembling a Poisson post-synaptic potential function:

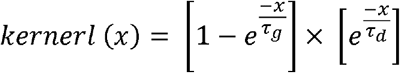

where *τ_g_* controls the rate of increase and *τ_d_* controls the rate of decay.

We further injected a small noise to the system by convolving the response kernel with Gaussian noise of *μ* = 0 and *σ*^2^ = 1. For each neuron, we multiplied the tuning curve and the convolved kernel to get the spatiotemporal firing rate for that neuron (**Fig S5D**). Saccade onsets were taken as the time when the normalized activity reached a fixed threshold of 1 unit.

For the simulated data with asymmetric bilateral inhibition, we followed the above steps (for both plans 1 and 2) until the activity for plan 1 reached y% of its peak response. This ‘y’ is given by an inhibition function (**Fig S5G**) which was constructed by dividing the neural response by the response in no-step trials. To simulate the inhibitory effect of plan 1 on plan 2, we temporally shifted the response for plan 2, to until after the time of first saccade onset (*t*_*s*1_), after plan 1 reached y% of its peak response and then interpolated the data in between using a cubic spline function.

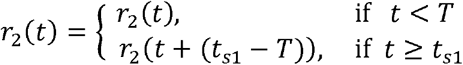

where *T* = *arg* (*y* × *max*(*r*_1_(*t*)))

To simulate the effect of plan 2 on plan 1, we first normalized plan 2’s response from 0 to 1, then multiplied it by the same inhibition factor, *y* and then subtracted it from plan 1’s response.

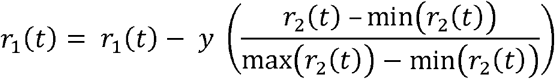

Therefore, the nature of inhibition was asymmetric and, in this way, plans 1 and 2 have the highest bilateral inhibition effect on each other for the shortest TSD and the least effect for the longest TSD. We estimated all the above hyper-parameters such that the simulated data closely resembled the experimental data.

For the simulated data with unilateral inhibition, we followed the same steps as above with the exception of the last step. That is, we modeled the effect of plan 1 on plan 2 by temporally shifting the plan 2 as described above but we did not account for the effect of plan 2 on plan 1.

After simulating the data, we followed the same data pre-processing step before dimensionality reduction similar to the experimental data.

#### Statistical testing

A two-sided Wilcoxon signed-rank test to analyze a single sample set of data. For group comparisons, the non-parametric Kruskal-Wallis test was used. Trials were considered to be independent observations as the TSDs on each trial was chosen randomly. All the results are presented as mean (± standard error of mean, SEM) and all tests are performed at a significance level of α = 0.05 unless otherwise mentioned.

## Data and code availability

All data is available in the main text or the supplementary materials. Raw data and codes are available upon reasonable request.

## Acknowledgments

We thank S. Sengupta for helping with behavioral training and Dr. A. Gopal P.A. for helping with data collection.

## Author Contributions

D.B. Performed research, Analyzed data, Wrote the paper. N.S. Analyzed data, Wrote the paper. A.M. Designed research, Wrote the paper.

---

**Figure S1.**
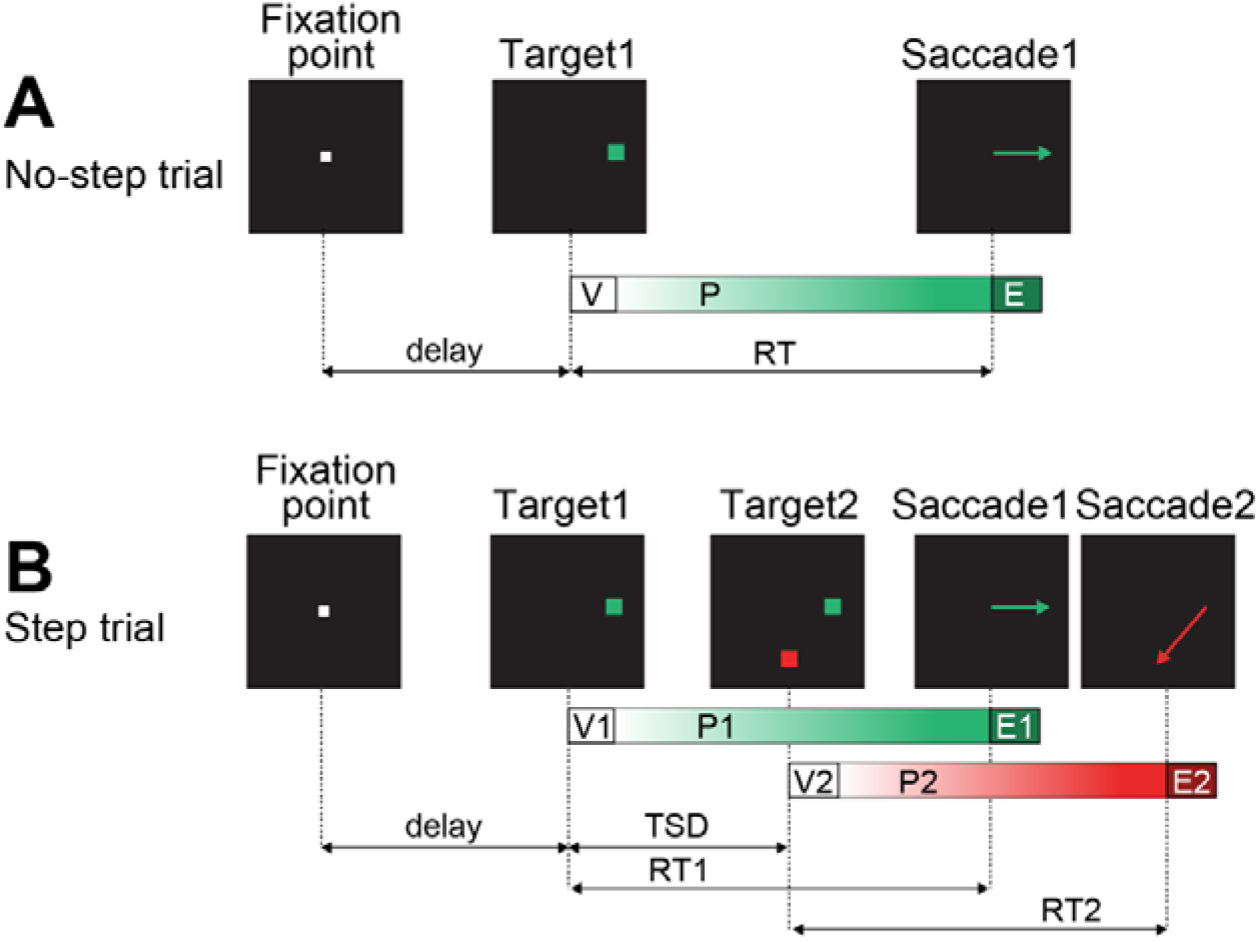
Schematic of the FOLLOW task (*related to Fig 1*) **A.** A representative no-step trial. The trial starts with the appearance of a central fixation point (FP), followed by the presentation of the green target (T1) at any one of the six possible peripheral locations. The monkey had to make a single saccade (S1) to the target to get a juice reward. In the representative framework, the processes leading to the culmination of a saccade are simplified to consist of three stages: visual encoding of stimuli(V), central planning (P), and saccade execution (E). RT refers to the reaction time. **B.** A representative step trial. Similar to no-step trials, a step trial started with central fixation, after which a green target (T1) appeared. A second red target (T2) was then presented after T1. A variable delay separated the first and the second target onsets (target step delay; TSD). The monkey had to make a sequence of two saccades (S1, S2) to the two targets in order of their appearance to get rewarded in step trials. The abbreviations used are the same as in **A**, but for two saccades.

**Figure S2.**
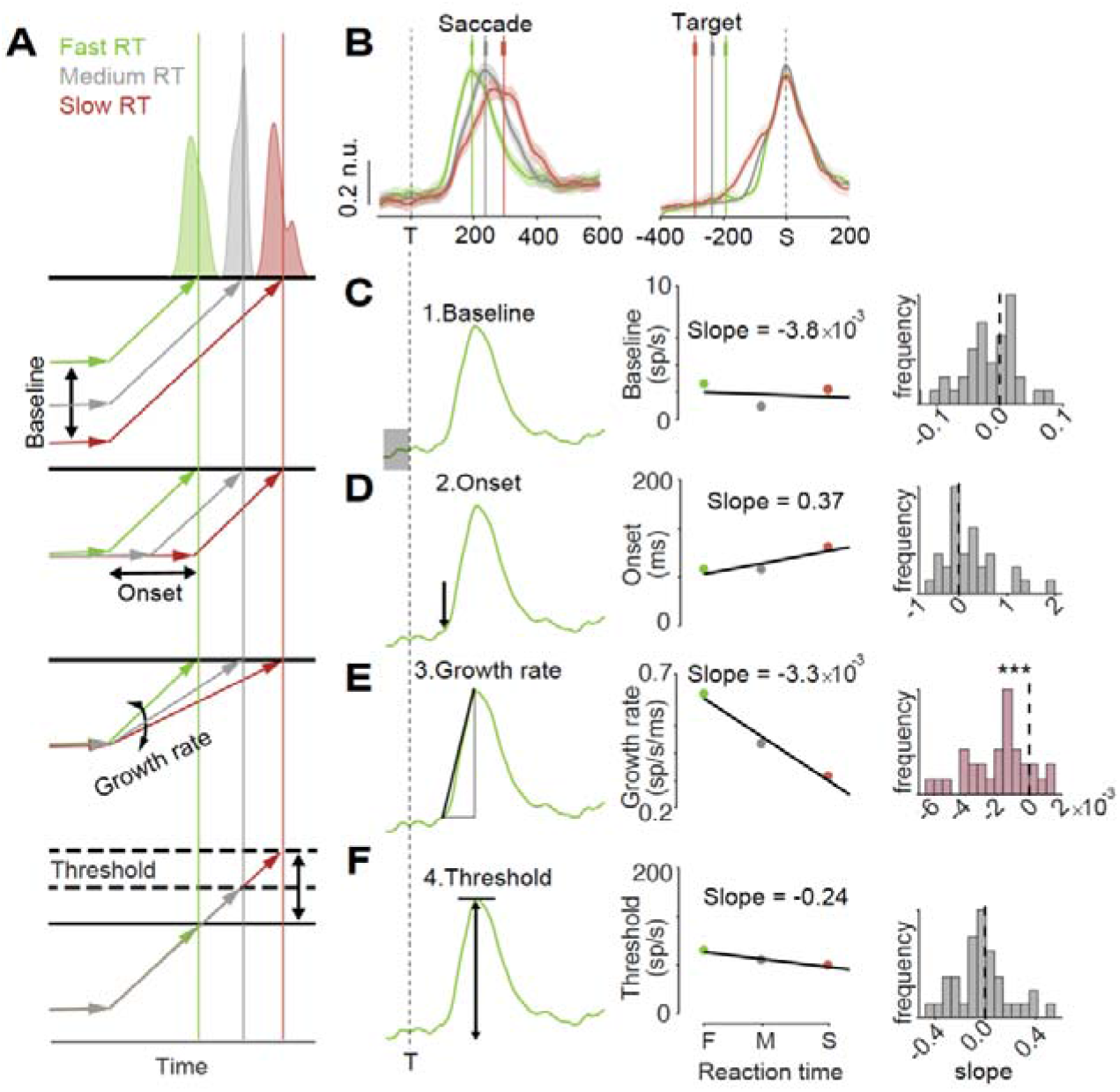
Population activity of FEF movement neurons in no-step trials (*related to Fig 3*) **A.** Schematic of possible adjustments of neural activity related to processing bottlenecks. The lengthening of the reaction time may be due to four possible adjustments according to accumulator dynamics. Each schematic shows a noise-free, simplistic accumulation process, starting from the baseline and reaching up to the threshold for saccade initiation. Representative reaction time distributions are plotted above each of the schematics. Lowering of baseline activity, delaying the onset of activity, slowing down the rate of growth, and increasing the threshold level can either singly or in combination, bring about increased reaction times. **B.** Left: FEF movement neuron population activity encoding for different reaction times (short: green, medium: gray, long: red), aligned to target onset (vertical broken line). Mean saccade onset times for all the reaction time conditions are shown as vertical colored lines with s.e.m error bars. Right: same as left but activity aligned to saccade onset (vertical broken line). Shading indicates mean ± SEM. **C.** Illustration of the measurement of accumulator parameter: baseline. Left: Schematic of a spike density function of a representative neuron aligned to target onset showing the baseline value. Middle: The baseline was calculated for short, medium and long reaction times for a representative neuron and the slope of the best fit was calculated. Right: Histogram of distribution of slopes measured this way for all the neurons was compared with zero (vertical broken line). **D.** Same as **C** but for measurement of onset. **E.** Same as **C** but for measurement of growth rate. **F.** Same as **C** but for measurement of threshold.

**Figure S3:**
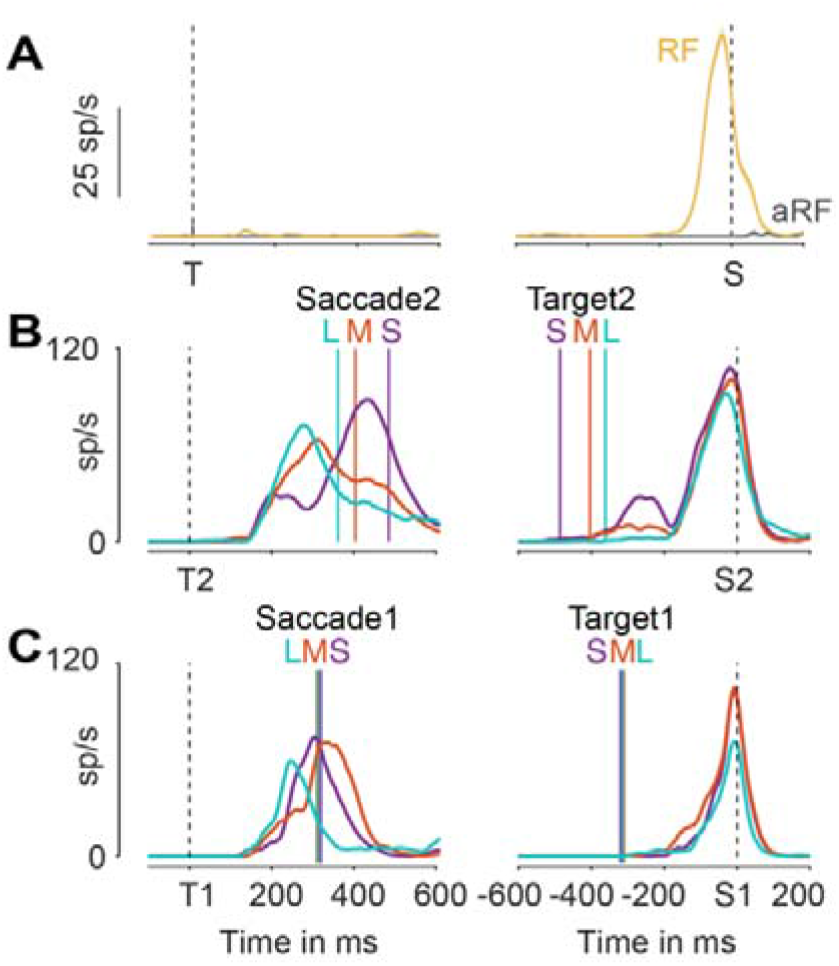
Single neuron example for processing bottlenecks *(related to Fig 3)* **A.** A representative FEF movement neuron aligned to target onset (T) and saccade onset (S) in a memory guided task for saccades into the RF and saccades out of RF. **B.** Left: Neural activity, a representative neuron encoding for different TSD conditions (short, medium, long TSD), aligned to target 2 onset. Right: same as left but activity aligned to S2 onset. **C.** Same as **B** but for neural activity aligned to target 1 (left) and saccade 1 (right).

**Figure S4:**
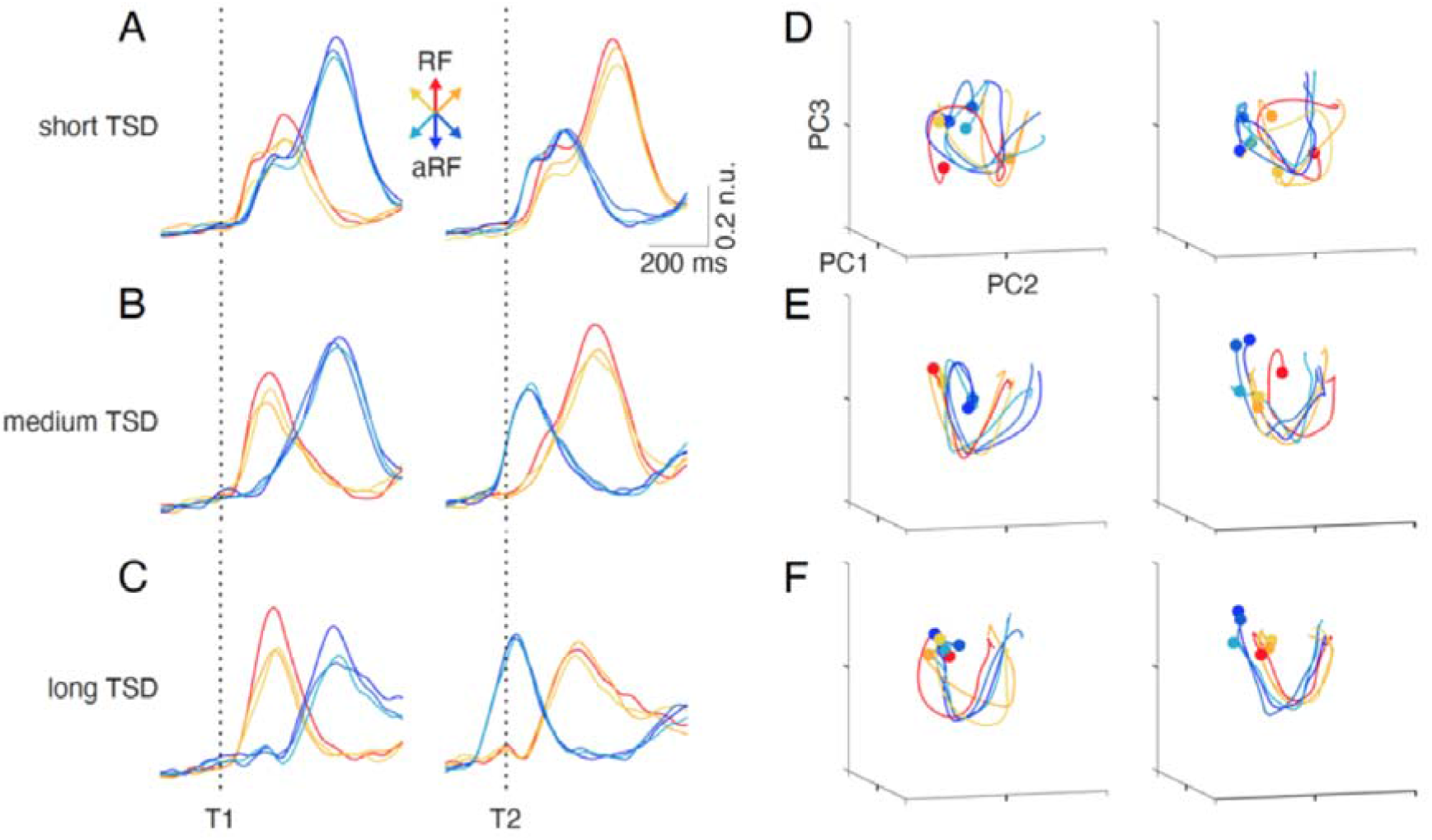
Population dynamics for all target positions (*related to Fig 4*) **A.** Soft-normalized mean population neural responses towards all six target positions (3 towards RF, shown in warm colors and 3 out of RF, shown in cold colors; see inset for colors), aligned to target 1 (left) and target 2 (right) for short TSD trials (n.u. = normalized unit). **B.** Same as **A**, but for medium TSD trials. **C.** Same as **A**, but for long TSD trials. **D.** First three PCs for each target position shown in **A**, plotted against each other for target 1 (left) and target 2 (right) related responses for short TSD trials. Filled circle markers indicate the starts of the respective trajectories. **E.** Same as **D**, but for medium TSD trials shown in **B**. **F.** Same as **D**, but for long TSD trials shown in **C**.

**Figure S5:**
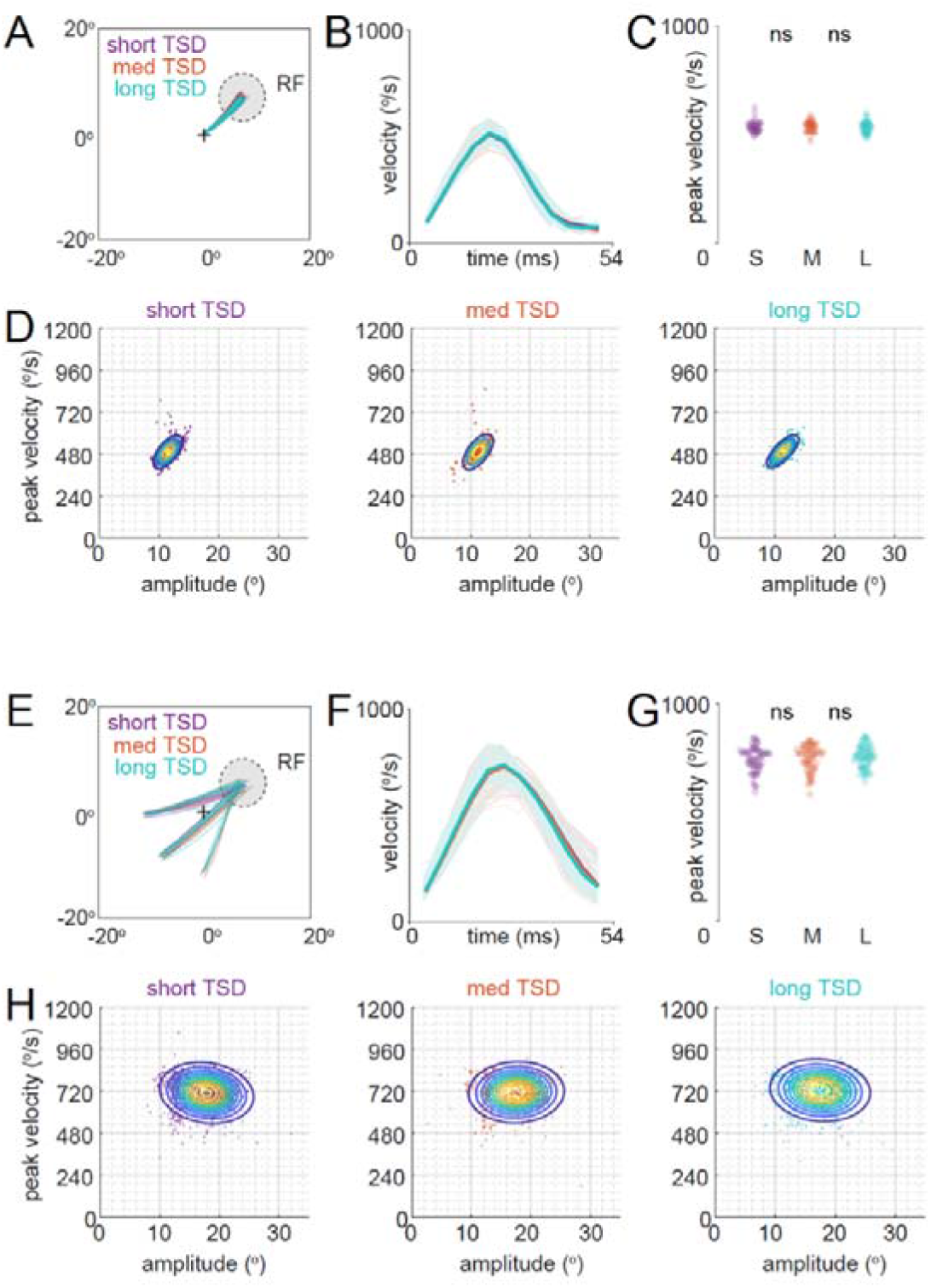
Saccade kinematics with different TSDs (*related to Fig 4*) **A.** Saccade 1 trajectories, towards RF (filled, broken circle), during short, medium and long TSD trials from a representative session. The central cross denotes the fixation point. **B.** Mean saccade 1 velocity profiles for short, medium and long TSD trials (thick lines) superimposed on saccade 1 velocity profiles from individual trials (thin lines). Shading indicates mean ± SEM. **C.** Peak saccade 1 velocities from each session for short, medium and long TSD trials. P values: short-medium: 0.81, ranksum test; medium-long: 0.60, ranksum test. **D.** Saccade 1 main sequence for short (left), medium (center) and long (right) TSD trials. The overlaid contours represent the density of data. **E.** Saccade 2 trajectories, towards RF (filled, broken circle), during short, medium and long TSD trials from the same representative session as **A**. **F.** Same as **B**, but for saccade 2. **G.** Same as **C**, but for saccade 2. P values: short-medium: 0.92, ranksum test; medium-long: 0.80, ranksum test. **H.** Same as **D**, but for saccade 2.

**Figure S6:**
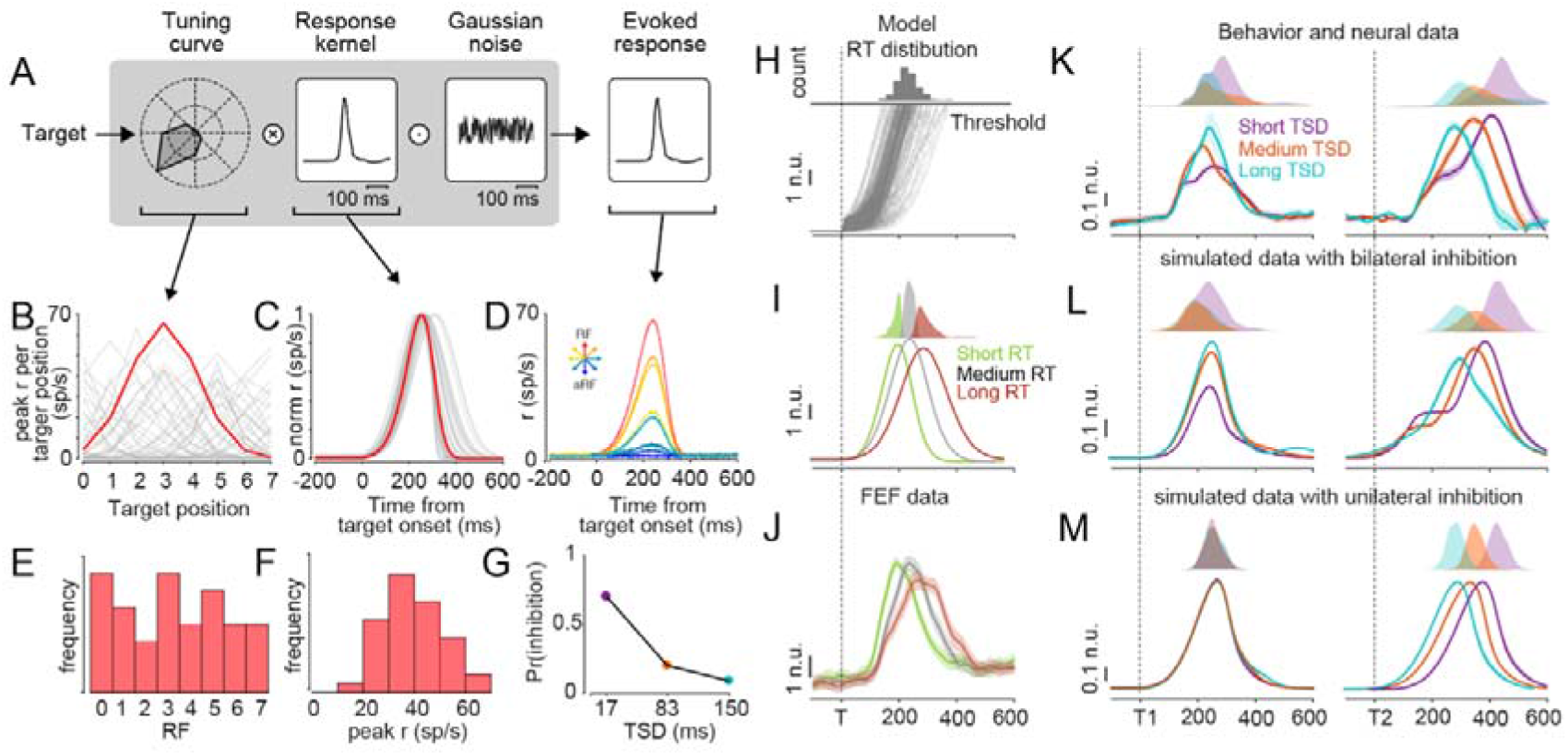
Data simulation process, reaction time and neural activity profiles (*related to Fig 5*) **A.** Schematic illustration of the simulation. **B.** Tuning curves (peak firing rates as a function of target positions) for all the simulated neurons. One representative neuron’s tuning curve is highlighted in purple. **C.** Response kernels (activity as a function of time) for all the simulated neurons. Same representative neuron’s stimulus kernel is highlighted in purple. **D.** Multiplication of tuning curves and stimulus kernels and addition of noise results in the simulated neuron’s activity for each of the 8 target positions. Activity towards the 8 target positions is shown for the same representative neuron as above. **E.** Distribution of RF positions (argmax(tuning curve)) of the simulated neurons shows a uniform spread of RFs across the simulated neurons. **F.** Distribution of peak firing rate (max(tuning curve)) of the simulated neurons shows a unimodal distribution with a mean ~ 37 sp/s. **G**. Probability of inhibition as a function of TSD **H**. Bottom: Example simulation for one session, for RF position, following the simulation pipeline shown in A. Top: reaction time distribution obtained from the simulation. **I.** Top: The same reaction time distribution from H, but uniformly divided into fast, medium and slow reaction times. Bottom: Average neural activity from the simulation in H, divided into fast, medium and slow reaction time conditions as explained above. **J.** Same FEF neural data from **Fig S2B** (divided into fast, medium and slow reaction time). **K.** Top: reaction time distributions of two monkeys for short, medium and long TSD trials for plan 1 (left) and plan 2 (right). Bottom: Same as **Fig 3C;** right: Same as **Fig 3A.** **L.** Top: reaction time distributions from the simulations with bilateral inhibition in the same format as above. Bottom: Same as **Fig 5F.** **M.** Top: reaction time distributions from the simulations with unilateral inhibition in the same format as above. Bottom: Same as **Fig 5B.**

## References

Alexander, G.E., DeLong, M.R., and Strick, P.L. (1986). Parallel organization of functionally segregated circuits linking basal ganglia and cortex. Annual review of neuroscience 9, 357–381.

Arnell, K.M., and Duncan, J. (2002). Separate and shared sources of dual-task cost in stimulus identification and response selection. Cognitive psychology 44, 105–147.

Basu, D., and Murthy, A. (2020). Parallel programming of saccades in the macaque frontal eye field: are sequential motor plans coactivated? Journal of neurophysiology 123, 107–119.

Becker, W., and Jürgens, R. (1979). An analysis of the saccadic system by means of double step stimuli. Vision research 19, 967–983.

Bhutani, N., Ray, S., and Murthy, A. (2012). Is saccade averaging determined by visual processing or movement planning? Journal of neurophysiology 108, 3161–3171.

Bhutani, N., Sureshbabu, R., Farooqui, A.A., Behari, M., Goyal, V., and Murthy, A. (2013). Queuing of concurrent movement plans by basal ganglia. Journal of Neuroscience 33, 9985–9997.

Bogacz, R., Brown, E., Moehlis, J., Holmes, P., and Cohen, J.D. (2006). The physics of optimal decision making: a formal analysis of models of performance in two-alternative forced-choice tasks. Psychological review 113, 700.

Boucher, L., Palmeri, T.J., Logan, G.D., and Schall, J.D. (2007). Inhibitory control in mind and brain: an interactive race model of countermanding saccades. Psychological review 114, 376.

Broadbent, D.E. (1971). Decision and stress.

Bruce, C.J., and Goldberg, M.E. (1985). Primate frontal eye fields. I. Single neurons discharging before saccades. Journal of neurophysiology 53, 603–635.

Churchland, M.M., Cunningham, J.P., Kaufman, M.T., Foster, J.D., Nuyujukian, P., Ryu, S.I., and Shenoy, K.V. (2012). Neural population dynamics during reaching. Nature 487, 51–56.

Coëffé, C., and O’regan, J.K. (1987). Reducing the influence of non-target stimuli on saccade accuracy: Predictability and latency effects. Vision research 27, 227–240.

Ding, L., and Gold, J.I. (2012). Separate, causal roles of the caudate in saccadic choice and execution in a perceptual decision task. Neuron 75, 865–874.

Elsayed, G.F., Lara, A.H., Kaufman, M.T., Churchland, M.M., and Cunningham, J.P. (2016). Reorganization between preparatory and movement population responses in motor cortex. Nature communications 7, 1–15.

Findlay, J.M. (1982). Global visual processing for saccadic eye movements. Vision research 22, 1033–1045.

Gold, J.I., and Shadlen, M.N. (2007). The neural basis of decision making. Annual review of neuroscience 30.

Goldman-Rakic, P.S., and Porrino, L.J. (1985). The primate mediodorsal (MD) nucleus and its projection to the frontal lobe. Journal of Comparative Neurology 242, 535–560.

Gopher, D., and Navon, D. (1980). How is performance limited: Testing the notion of central capacity. Acta psychologica 46, 161–180.

Hanes, D.P., Patterson, W.F., and Schall, J.D. (1998). Role of frontal eye fields in countermanding saccades: visual, movement, and fixation activity. Journal of neurophysiology 79, 817–834.

Hanes, D.P., and Schall, J.D. (1996). Neural control of voluntary movement initiation. Science 274, 427–430.

Heitz, R.P., and Schall, J.D. (2012). Neural mechanisms of speed-accuracy tradeoff. Neuron 76, 616–628.

Hikosaka, O., Takikawa, Y., and Kawagoe, R. (2000). Role of the basal ganglia in the control of purposive saccadic eye movements. Physiological reviews 80, 953–978.

Huerta, M.F., Krubitzer, L.A., and Kaas, J.H. (1986). Frontal eye field as defined by intracortical microstimulation in squirrel monkeys, owl monkeys, and macaque monkeys: I. Subcortical connections. Journal of Comparative Neurology 253, 415–439.

Jagadisan, U.K., and Gandhi, N.J. (2016). Disruption of fixation reveals latent sensorimotor processes in the superior colliculus. Journal of Neuroscience 36, 6129–6140.

Kahneman, D. (1973). Attention and effort, Vol 1063 (Citeseer).

Langer, T.P., and Kaneko, C.R. (1990). Brainstem afferents to the oculomotor omnipause neurons in monkey. Journal of Comparative Neurology 295, 413–427.

Lo, C.-C., and Wang, X.-J. (2006). Cortico–basal ganglia circuit mechanism for a decision threshold in reaction time tasks. Nature neuroscience 9, 956–963.

Markram, H., Toledo-Rodriguez, M., Wang, Y., Gupta, A., Silberberg, G., and Wu, C. (2004). Interneurons of the neocortical inhibitory system. Nature reviews neuroscience 5, 793–807.

Marois, R., and Ivanoff, J. (2005). Capacity limits of information processing in the brain. Trends in cognitive sciences 9, 296–305.

McLeod, P. (1977). A dual task response modality effect: Support for multiprocessor models of attention. Quarterly Journal of Experimental Psychology 29, 651–667.

McPeek, R.M., Han, J.H., and Keller, E.L. (2003). Competition between saccade goals in the superior colliculus produces saccade curvature. Journal of neurophysiology 89, 2577–2590.

McPeek, R.M., and Keller, E.L. (2002). Superior colliculus activity related to concurrent processing of saccade goals in a visual search task. Journal of Neurophysiology 87, 1805–1815.

McPeek, R.M., Skavenski, A.A., and Nakayama, K. (2000). Concurrent processing of saccades in visual search. Vision research 40, 2499–2516.

Middleton, F.A., and Strick, P.L. (2000). Basal ganglia and cerebellar loops: motor and cognitive circuits. Brain research reviews 31, 236–250.

Minken, A., Van Opstal, A., and Van Gisbergen, J. (1993). Three-dimensional analysis of strongly curved saccades elicited by double-step stimuli. Experimental Brain Research 93, 521–533.

Murthy, A., Ray, S., Shorter, S.M., Priddy, E.G., Schall, J.D., and Thompson, K.G. (2007). Frontal eye field contributions to rapid corrective saccades. Journal of Neurophysiology 97, 1457–1469.

Murthy, A., Ray, S., Shorter, S.M., Schall, J.D., and Thompson, K.G. (2009). Neural control of visual search by frontal eye field: effects of unexpected target displacement on visual selection and saccade preparation. Journal of Neurophysiology 101, 2485–2506.

Navon, D., and Miller, J. (2002). Queuing or sharing? A critical evaluation of the single-bottleneck notion. Cognitive psychology 44, 193–251.

Nosofsky, R.M., and Palmeri, T.J. (1997). Comparing exemplar-retrieval and decision-bound models of speeded perceptual classification. Perception & Psychophysics 59, 1027–1048.

Parent, A., and Hazrati, L.-N. (1995a). Functional anatomy of the basal ganglia. I. The cortico-basal ganglia-thalamo-cortical loop. Brain research reviews 20, 91–127.

Parent, A., and Hazrati, L.-N. (1995b). Functional anatomy of the basal ganglia. II. The place of subthalamic nucleus and external pallidium in basal ganglia circuitry. Brain research reviews 20, 128–154.

Pashler, H. (1994). Dual-task interference in simple tasks: data and theory. Psychological bulletin 116, 220.

Phillips, A.N., and Segraves, M.A. (2010). Predictive activity in macaque frontal eye field neurons during natural scene searching. Journal of neurophysiology 103, 1238–1252.

Port, N.L., and Wurtz, R.H. (2003). Sequential activity of simultaneously recorded neurons in the superior colliculus during curved saccades. Journal of neurophysiology 90, 1887–1903.

Purcell, B.A., Heitz, R.P., Cohen, J.Y., and Schall, J.D. (2012). Response variability of frontal eye field neurons modulates with sensory input and saccade preparation but not visual search salience. Journal of neurophysiology 108, 2737–2750.

Purcell, B.A., Heitz, R.P., Cohen, J.Y., Schall, J.D., Logan, G.D., and Palmeri, T.J. (2010). Neurally constrained modeling of perceptual decision making. Psychological review 117, 1113.

Ramakrishnan, A., and Murthy, A. (2013). Brain mechanisms controlling decision making and motor planning. In Progress in brain research (Elsevier), pp. 321–345.

Ratcliff, R., Hasegawa, Y.T., Hasegawa, R.P., Smith, P.L., and Segraves, M.A. (2007). Dual diffusion model for single-cell recording data from the superior colliculus in a brightness-discrimination task. Journal of neurophysiology 97, 1756–1774.

Ratcliff, R., and Rouder, J.N. (1998). Modeling response times for two-choice decisions. Psychological science 9, 347–356.

Ratcliff, R., and Smith, P.L. (2004). A comparison of sequential sampling models for two-choice reaction time. Psychological review 111, 333.

Ray, S., Bhutani, N., and Murthy, A. (2012). Mutual inhibition and capacity sharing during parallel preparation of serial eye movements. Journal of vision 12, 17–17.

Ray, S., Pouget, P., and Schall, J.D. (2009). Functional distinction between visuomovement and movement neurons in macaque frontal eye field during saccade countermanding. Journal of Neurophysiology 102, 3091–3100.

Ray, S., Schall, J.D., and Murthy, A. (2004). Programming of double-step saccade sequences: modulation by cognitive control. Vision research 44, 2707–2718.

Ruthruff, E., Pashler, H.E., and Klaassen, A. (2001). Processing bottlenecks in dual-task performance: Structural limitation or strategic postponement? Psychonomic bulletin & review 8, 73–80.

Sato, T., Murthy, A., Thompson, K.G., and Schall, J.D. (2001). Search efficiency but not response interference affects visual selection in frontal eye field. Neuron 30, 583–591.

Segraves, M.A. (1992). Activity of monkey frontal eye field neurons projecting to oculomotor regions of the pons. Journal of Neurophysiology 68, 1967–1985.

Segraves, M.A., and Goldberg, M.E. (1987). Functional properties of corticotectal neurons in the monkey’s frontal eye field. Journal of Neurophysiology 58, 1387–1419.

Sendhilnathan, N., Basu, D., Goldberg, M.E., Schall, J.D., and Murthy, A. (2021). Neural correlates of goal-directed and non–goal-directed movements. Proceedings of the National Academy of Sciences 118.

Sendhilnathan, N., Basu, D., and Murthy, A. (2017). Simultaneous analysis of the LFP and spiking activity reveals essential components of a visuomotor transformation in the frontal eye field. Proceedings of the National Academy of Sciences 114, 6370–6375.

Sendhilnathan, N., Basu, D., and Murthy, A. (2020). Assessing within-trial and across-trial neural variability in macaque frontal eye fields and their relation to behaviour. European Journal of Neuroscience.

Sharika, K., Ramakrishnan, A., and Murthy, A. (2008). Control of predictive error correction during a saccadic double-step task. Journal of neurophysiology 100, 2757–2770.

Shen, K., and Paré, M. (2014). Predictive saccade target selection in superior colliculus during visual search. Journal of Neuroscience 34, 5640–5648.

Sigman, M., and Dehaene, S. (2005). Parsing a cognitive task: a characterization of the mind’s bottleneck. PLoS biology 3, e37.

Smith, P.L., and Van Zandt, T. (2000). Time-dependent Poisson counter models of response latency in simple judgment. British Journal of Mathematical and Statistical Psychology 53, 293–315.

Sommer, M.A., and Wurtz, R.H. (2000). Composition and topographic organization of signals sent from the frontal eye field to the superior colliculus. Journal of Neurophysiology 83, 1979–2001.

Somogyi, P. (1977). A specific ‘axo-axonal’interneuron in the visual cortex of the rat. Brain Res 136, 345–350.

Thompson, K.G., Bichot, N.P., and Schall, J.D. (1997). Dissociation of visual discrimination from saccade programming in macaque frontal eye field. Journal of neurophysiology 77, 1046–1050.

Thompson, K.G., Hanes, D.P., Bichot, N.P., and Schall, J.D. (1996). Perceptual and motor processing stages identified in the activity of macaque frontal eye field neurons during visual search. Journal of neurophysiology 76, 4040–4055.

Tian, J., Schlag, J., and Schlag-Rey, M. (2000). Testing quasi-visual neurons in the monkey’s frontal eye field with the triple-step paradigm. Experimental brain research 130, 433–440.

Tombu, M., and Jolicœur, P. (2003). A central capacity sharing model of dual-task performance. Journal of Experimental Psychology: Human Perception and Performance 29, 3.

Usher, M., and McClelland, J.L. (2001). The time course of perceptual choice: the leaky, competing accumulator model. Psychological review 108, 550.

Viviani, P., and Swensson, R.G. (1982). Saccadic eye movements to peripherally discriminated visual targets. Journal of Experimental Psychology: Human Perception and Performance 8, 113.

Welford, A. (1967). Single-channel operation in the brain. Acta psychologica 27, 5–22.

Welford, A.T. (1952). The psychological refractory period and the timing of high-speed performance-a review and a theory. British Journal of Psychology 43, 2.

Westheimer, G. (1954). Eye movement responses to a horizontally moving visual stimulus. AMA archives of ophthalmology 52, 932–941.

Wheeless, L.L., Boynton, R.M., and Cohen, G.H. (1966). Eye-movement responses to step and pulse-step stimuli. JOSA 56, 956–960.

Woodman, G.F., Kang, M.-S., Thompson, K., and Schall, J.D. (2008). The effect of visual search efficiency on response preparation: neurophysiological evidence for discrete flow. Psychological science 19, 128–136.

Wu, E.X., Gilani, S.O., van Boxtel, J.J., Amihai, I., Chua, F.K., and Yen, S.-C. (2013). Parallel programming of saccades during natural scene viewing: Evidence from eye movement positions. Journal of vision 13, 17–17.

Wurtz, R.H., and Hikosaka, O. (1986). Role of the basal ganglia in the initiation of saccadic eye movements. In Progress in brain research (Elsevier), pp. 175–190.

Zambarbieri, D., Schmid, R., and Ventre, J. (1987). Saccadic eye movements to predictable visual and auditory targets. In Eye Movements from Physiology to Cognition (Elsevier), pp. 131–140.

Zylberberg, A., Ouellette, B., Sigman, M., and Roelfsema, P.R. (2012). Decision making during the psychological refractory period. Current biology 22, 1795–1799.

